# Kinetic sequencing (*k*-Seq) as a massively parallel assay for ribozyme kinetics: utility and critical parameters

**DOI:** 10.1101/2020.12.02.407346

**Authors:** Yuning Shen, Abe Pressman, Evan Janzen, Irene Chen

## Abstract

Characterization of genotype-phenotype relationships of genetically encoded molecules (e.g., ribozymes) requires accurate quantification of activity for a large set of molecules. Kinetic measurement using high-throughput sequencing (e.g., *k*-Seq) is an emerging assay applicable in various domains that potentially scales up measurement throughput to 10^5^ ~ 10^6^ unique sequences. However, technical challenges introduced by sequence heterogeneity and DNA sequencing must be understood to realize the utility and limitations of such assays. We characterized the *k*-Seq method in terms of model identifiability, effects of sequencing error, accuracy and precision using simulated datasets and experimental data from a variant pool constructed from previously identified ribozymes. Relative abundance, kinetic coefficients, and measurement noise were found to affect the measurement of each sequence. We introduced bootstrapping to robustly quantify the uncertainty in estimating model parameters and proposed interpretable metrics to quantify model identifiability. These efforts enabled the rigorous reporting of data quality for individual sequences in *k*-Seq experiments. Critical experimental factors were examined, and general guidelines are proposed to maximize the number of sequences having precisely estimated and identifiable kinetic coefficients from *k*-Seq data. Practices analogous to those laid out here could be applied to improve the rigor of similar sequencing-based assays.

## INTRODUCTION

Determining the genotype-phenotype relationships for any large set of biomolecules requires a high-throughput method to measure the chemical activity of each sequence in the set. For catalytic nucleic acids, measuring the activity of each unique sequence in a diverse population is an important goal. Ideally, methods to accomplish this would: a) yield accurate activity measurements for individual sequences, b) achieve high throughput, such as by parallelization, to cover a large number of variants in sequence space, and c) be adaptable to different ribozymes (and deoxyribozymes).

High-throughput sequencing (HTS) provides an opportunity to address these goals. The large amount of sequence data should allow high accuracy count data for many sequences in a high throughput, parallelized format. In principle, if reacted and unreacted molecules can be separated from each other and sequenced independently, HTS could quantify the extent of a reaction for millions of different sequences simultaneously. Since nucleic acid sequences act as their own ‘barcodes’, use of sequencing as the assay would avoid the need to isolate and test each unique sequence individually. Such a method can measure the activity of each sequence in a population of functional molecules, at multiple time points, substrate concentrations, or other variable conditions. HTS-based kinetic measurements have been proposed and demonstrated with nucleic acids, including catalytic DNA (1), catalytic RNA (2–5), substrate RNA (“HTS-Kin”) (6), RNA aptamers (7), and transcription factors (TF) binding to DNA (8). In these studies, approximately 10^3^ ~ 10^6^ unique sequences are measured, depending on the experimental design. Similar approaches have also been developed for proteins, notably an assay of ligand binding affinities through mRNA display (9), ‘deep mutational scanning’, in which the phenotype of fitness is assayed for many mutants by deep sequencing (10), and a large-scale measurement of dose-response curves (11). However, the development of these massively parallel measurements also raises important questions about experimental design, normalization, sequencing errors, and measurement accuracy and precision. To date, there is a relative lack of critical discussion addressing the theoretical and experimental effects of such variables on the outcome of high-throughput measurements. The purpose of the present work is to examine these issues and develop appropriate methodology to address them in the context of assaying ribozymes.

Here we focus on kinetic sequencing (*k*-Seq), a recently reported method for quantification of kinetics in a mixed pool of sequences (2). Advantages of *k*-Seq include absolute, rather than relative (5, 6) measurements, as well as the lack of requirement for specialized instrumentation (4, 7). A general schema of *k*-Seq is described as follows (Figure 1): an input pool is designed containing sequences of interest (e.g., candidate ribozymes). Aliquots of the input pool are reacted under different experimental conditions, such as different substrate concentrations or different time points. Then, reacted and unreacted molecules are separated. Each pool is converted to a DNA library and prepared for sequencing. Absolute measurement of reacted (or unreacted) quantities is also performed to allow normalization. Reads generated from HTS are subjected to quality control and de-replicated to generate a ‘count’ table of the copy number for each sequence detected in a sample. Count data are normalized to absolute abundance and fit to the appropriate kinetic model to estimate the rate constants and other parameters of interest.

**Figure 1:**
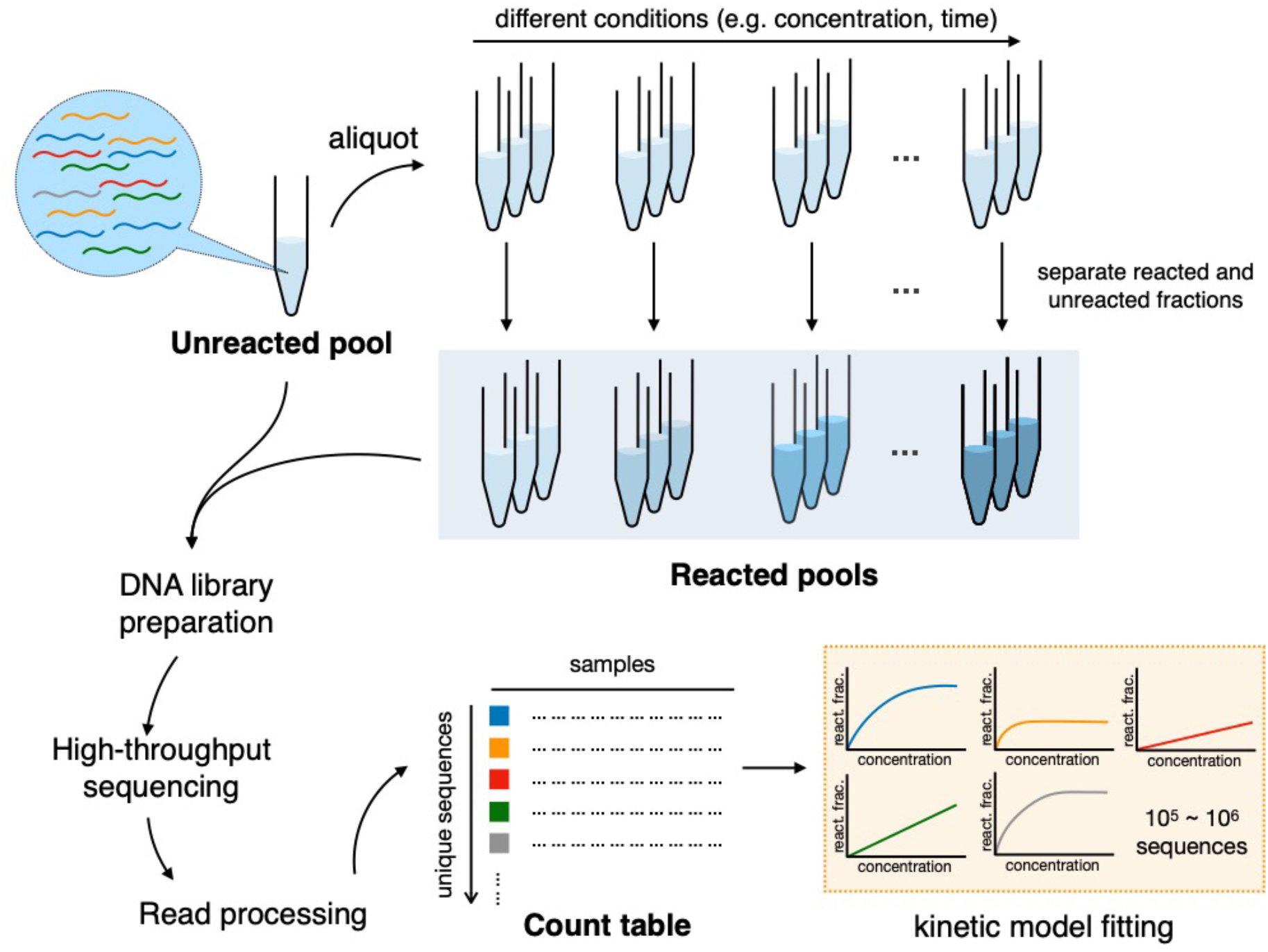
General scheme of *k*-Seq experiment and analysis. A heterogeneous input pool containing nucleic acids is reacted at different experimental conditions (e.g., different substrate concentrations or different reaction time). Reacted and unreacted molecules are separated and either (or both) of these fractions is prepared for high-throughput sequencing. The reads from DNA sequencing are processed to obtain a count table for each unique sequence across samples, normalized by a standard, and abundances across samples are fit into a kinetic model to estimate parameters (e.g., rate constants). react. frac. = reacted fraction.

Multiple issues could potentially limit the applicability of *k*-Seq and related methods. Kinetic measurements require properly chosen experimental conditions (e.g., substrate concentrations or time points) for a sufficient dynamic range. For a heterogeneous pool where different sequences have different optimal conditions, the conditions chosen will represent a compromise for some sequences. For example, in a single time-point (reacted vs. unreacted) experiment determining enzyme kinetics over 4096 RNA substrates (6), the choice of reaction time would be optimal for either highly active RNA substrates or less active ones, but not all. Previous work determining kinetics for ribozymes by *k*-Seq also showed that characterization of less active ribozymes was limited when the kinetic model parameters (rate constant *k* and maximum amplitude of reaction *A*) could not be independently estimated due to model identifiability problems (2). While the number of experimental conditions could be increased to address this problem, with HTS this potential solution would quickly become prohibitively expensive in time and resources. Therefore, it is important to rigorously understand how the choice of conditions would affect the estimation of kinetic parameters and trustworthiness of measurements on each sequence.

Another consideration unique to HTS-based kinetic measurements derives from the inevitability of sequencing errors in the data. Sequencing error might misidentify a molecule as a nearby sequence variant, and subsequently change the quantification of both the true and incorrect sequences. This could be particularly problematic when one sequence is present in high abundance relative to others, and thus creates a relatively large number of misidentified reads that confound the quantitation of related sequences. It is important to note that this problem cannot be solved by increasing the number of replicates or sequencing depth, since the number of erroneous reads rises in proportion to the number of reads (a systematic bias rather than random noise).

A final concern is assessing the accuracy and precision of *k*-Seq measurements. Using discrete count data (number of reads of a particular sequence) as the approximation of a sequence’s relative abundance in the sample introduces some complexity in assessing measurement accuracy, particularly at low counts where large stochastic variation exists. One approach taken in earlier works is to limit the library size to thousands of sequences which are each present in high copy number (high coverage of sequences) (1, 6). However, this requirement might unnecessarily restrict the applicability of *k*-Seq and related methods. When extending the approach to larger libraries or libraries with uneven coverage (e.g., a ‘doped’ variant pool or pool obtained from *in vitro* selection), high error may be associated with measurements of sequences with low counts (3, 4, 9). While one may simply exclude sequences with counts lower than a cutoff value, it is not obvious how to choose such cutoffs. Instead, it would be desirable to estimate the uncertainty (e.g., confidence intervals) on fitted parameter values for each sequence, given the experimental scenario of a low number of replicates. This approach would maximize the return of accurate information from a *k*-Seq experiment.

In this work, we performed a *k*-Seq experiment on a newly designed pool of variants based on ribozymes previously isolated from *in vitro* selection (2). These ribozymes react with an activated amino acid substrate (biotinyl-Tyr(Me)-oxazolone, or BYO) to produce aminoacyl-RNA, and are characterized by pseudo-first order kinetics. Coupled with theoretical and simulation studies, we systematically characterize model identifiability, accuracy, and precision of estimation from HTS data. We discuss key factors to consider when optimizing experimental design for *k*-Seq experiments. Lastly, we present a Python package of analysis tools for users undertaking *k*-Seq experiments. Incorporation of these techniques and lessons into HTS-based kinetic measurements should improve rigorous quantitative inference from similar experiments.

## MATERIALS AND METHODS

### Pseudo-first order kinetic model

As previously described in (2), we used a pseudo-first order rate equation to model the fraction of ribozymes reacted with substrate BYO:

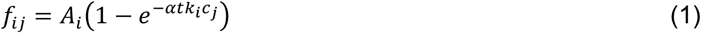

where *f*_*ij*_ is the reacted fraction for sequence *i* in sample *j* with initial BYO concentration *c*_*j*_. The kinetics for sequence *i* is characterized by *A*_*i*_ (the maximum amplitude of reaction) and *k*_*i*_ (the rate constant). BYO is present in vast excess compared to ribozymes in the experiments. A constant reaction time *t* = 90 min was used, corresponding to a coefficient *α* = 0.479 that accounts for the degradation of BYO during the reaction, as measured in (2). The product *k*_*i*_*A*_*i*_ was used as a combined parameter for activity measure and represents the initial rate of the reaction.

### Model identifiability for different *k*_*i*_ and *A*_*i*_

Depending on the BYO concentrations, the pseudo-first order kinetic model may be characterized as practically unidentifiable (12) where *k*_*i*_ and *A*_*i*_ are sensitive to noise and cannot be separately estimated. We evaluated this effect for each sequence using bootstrapping (resampling data with replacement; see below for details). We designed two metrics to score the model identifiability from bootstrapping results: *σ*_*A*_ and 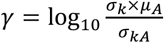, where *σ*_*A*_, *σ*_*k*_, *σ*_*kA*_ are the standard deviations for *A*, *k*, and *kA*, and *μ*_*A*_ is the mean value of *A*, from bootstrapped samples for each sequence. The score *γ* is the ratio of standard deviations for the rate constant *k* over the combined parameter *kA*, scaled by estimated *A*. If *k* and *A* are well-estimated independently, the ratio would be close to 1 and *γ* would be close to 0; if *k* and *A* are not estimated independently such that *k* has larger variance than *kA* in estimation, *γ* would be larger than 0. While bootstrapping can capture the noise in the experimental measurements through resampling, we also separately examined the convergence of fitting through 20 independent runs with random initial values of *k* and *A* sampled from Uniform(0, 10) min^−1^M^−1^ and Uniform(0, 1) respectively, using the original data (no resampling). The convergence of fitting was evaluated by the range of fitted *A* values (Δ*A* = *A*_max_ – *A*_min_) and used as a third candidate metric for model identifiability (higher Δ*A* ~ less identifiable).

To study model identifiability with controlled variables, we created a simulated reacted fraction dataset containing 10,201 (101^2^) sequences with the *log*_10_ of true values of *k* and *A* on a 101-by-101 grid across the region with log_10_ *k* ∈ [−1, 3] and log_10_ *A* ∈ [−2, 0]. The true reacted fraction 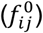 for sequence *i* in sample *j* with initial BYO concentration *c*_*j*_ was calculated based on the pseudo-first order rate equation (Equation 1). To model the effect of measurement noise, we added a Gaussian error term err_ij_ on the true reacted fraction 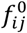 with the variance equal to 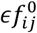, where the *∊* is the relative error of the reacted fraction:

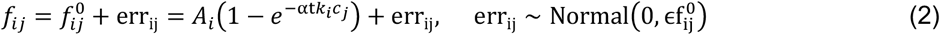

where *f*_*ij*_ is the observed reacted fraction. We chose *∊* = 0, 0.2, 0.5, 1.0 to evaluate the effect of measurement error. Negative values of *f*_*ij*_ were reassigned to be zero. These simulated reacted fractions were then used to estimate *k*_*i*_ and *A*_*i*_ for each simulated sequence using least-squares fitting as described below.

To study whether experimental and sequencing effort would be best spent extending the substrate concentration range vs. performing additional replicates, we simulated reacted fraction data for three different sets of BYO concentrations: *i*) standard set: 2, 10, 50, and 250 μM with triplicates, for 12 samples in total, as done previously to analyze a pool after *in vitro* selection (2); *ii*) additional replicates: 2, 10, 50, and 250 μM with four replicates each, for 16 samples in total; and *iii*) extended substrate range: 2, 10, 50, 250, and 1250 μM with triplicates, for 15 samples in total, as done in the variant pool experiment reported here.

### Sequence variant pool for aminoacylation assay

Four DNA libraries were obtained from Keck Biotechnology Laboratory, with the sequence 5′-<u>GATAATACGACTCACTATA</u>GGGAATGGATCCACATCTACGAATTC-[central variable region, length 21]-TTCACTGCAGACTTGACGAAGCTG-3′ (nucleotides upstream of the transcription start site are underlined). For each library, the central region corresponded to a 21-nucleotide variable sequence, based on a ribozyme family wild-type sequence (S-1A.1-a, S-1B.1-a, S-2.1-a, or S-3.1-a previously identified in (2), see Table S1 for sequences), with partial randomization at each position (specified to be 91% of the wild-type nucleotide and 3% of each base substitution at all 21 positions, i.e., a ‘doped’ pool). RNA was transcribed using HiScribe T7 RNA polymerase (New England Biolabs) and purified by denaturing polyacrylamide gel electrophoresis (PAGE) as previously described (2). An equimolar mixture of these four RNA libraries (the variant pool) was prepared for the *k*-Seq experiment.

### *k*-Seq experiment on the mixed pool of variants

Reactions were carried out in triplicates at 2, 10, 50, 250, and 1250 μM BYO for 90 minutes, following the incubation, RNA recovery and reverse-transcription protocols used in (2). Briefly, in each 50 μL *k*-Seq reaction, 2 μg total RNA was reacted with BYO. The reactions were stopped by desalting and placed on ice. Reacted sequences were isolated by pull-down with Streptavidin MagneSphere paramagnetic beads (Promega) and eluted with formamide/EDTA. 10 % of eluted RNA was taken to measure the total RNA amount using Qubit and qPCR (see below). A ‘spike-in’ RNA was added as an alternative quantification method (see below). RNA was prepared for sequencing by reverse transcription and PCR (RT-PCR), with primers complementary to the fixed sequences flanking the variable region (GATAATACGACTCACTATAGGGAATGGATCCACATCTACGAATTC, forward; CAGCTTCGTCAAGTCTGCAGTGAA, reverse). DNA from each of 15 samples was barcoded and pooled together in equal proportions. A reverse-transcribed unreacted sample was added at three times the total amount of DNA of one reacted sample to have similar total sequencing depth with each set of BYO concentration triplicates. Pooled DNA was sequenced on an Illumina NextSeq 500 with 150 bp paired-end run (Biological Nanostructures Laboratory, California NanoSystems Institute at UCSB), using a high output reagent kit expected to produce > 400 million reads.

### Quantitation of total amount of RNA per sample

We used two methods to quantify the absolute amount of RNA in *k*-Seq samples. Method 1 measured the amount of RNA in the samples after elution using Qubit or qPCR. For reactions carried out at 250 and 1250 μM, 10% of the RNA recovered after elution was quantified with an Invitrogen Qubit 3.0 fluorometer. If the recovered RNA was below the limit of detection by Qubit, quantitation was done by reverse-transcription-qPCR (same PCR primers as in the previous section) using a Bio-Rad C1000 thermal cycler with CFX96 Real-Time PCR block, SuperScript III RTase (Invitrogen), Phusion® High-Fidelity Polymerase (New England Biolabs), and SYBR Green (0.5x) (Figure S1). Method 2 used an internal standard (spike-in sequence) and data were normalized using sequencing results. A control spike-in sequence (5′-<u>GATAATACGACTCACTATA</u>GGGAATGGATCCACATCTACGAATTC-AAAAACAAAAACAAAAACAAA-TTCACTGCAGACTTGACGAAGCTG-3′, promoter underlined) was added to each sample before reverse transcription. 0.04, 0.2, 1, 2, and 2 ng of spike-in RNA was added to samples with 2, 10, 50, 250, 1250 μM BYO concentration respectively. 10 ng of spike-in RNA was added to the unreacted pool sample. The total RNA recovered (*Q*_*j*_) in sample *j* was calculated as

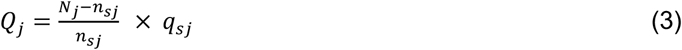

where *N*_*j*_ is the total number of reads in sample *j*, *n*_*sj*_ is the total reads of sequences within 2 edit distance (i.e., number of substitutions, insertions, or deletions) of the spike-in sequence in sample *j*, and *q*_*sj*_ is the quantity of spike-in sequence added to sample *j* after the reaction.

### Processing of *k*-Seq reads

FASTQ files of de-multiplexed paired-end Illumina reads were processed using EasyDIVER (13) to count the number of reads of each unique sequence in each sample. The forward and reverse reads were joined using pandaSeq (14) with the options ‘-a’ to join the paired-end reads before trimming and ‘completely_miss_the_point:0’ to enforce absolute matching in the overlapped variable region (any pairs with a disagreement between forward and reverse reads were discarded), thus minimizing sequencing errors. After joining, forward and reverse primers were trimmed by pandaSeq using ‘CTACGAATTC’ (forward) and ‘CTGCAGTGAA’ (reverse) adapter sequences. Next, multiple lanes for the same sample were combined and reads were de-replicated to give unique sequences and counts. The generated count files were analyzed using the ‘k-seq’ python package (https://github.com/ichen-lab-ucsb/k-seq). We collected all detected sequences in unreacted and/or reacted samples and discarded those that were not 21 nucleotides long as well as those within an edit distance of 2 from the spike-in sequence. The absolute amount (ng) for sequence *i* in sample *j* were quantified using total RNA recovered *Q*_*j*_ and number of reads:

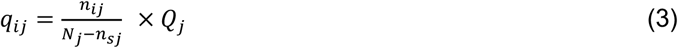

The reacted fractions for sequences in reacted samples were further calculated as the ratio to the absolute amount in the unreacted pool (*q*_0*j*_):

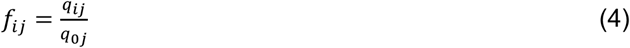

To be considered analyzable for fitting, a sequence needed at least one non-zero value among reacted samples as well as a non-zero count in the unreacted sample; non-analyzable sequences were discarded.

### Experimental coverage of mutants in the variant pool

Coverage of mutants in the variant RNA pool was analyzed from sequencing results of the unreacted pool. Unique sequences were classified into family centers (Hamming distance *d* = 0), single mutants (*d* = 1), double mutants (*d* = 2), triple mutants (*d* = 3), and others (*d* ≥ 4) based on their Hamming distance (number of substitutions) to the nearest family center (S-2.1-a, S-1A.1-a, S-1B.1-a, or S-3.1-a). The coverage fraction for a certain class of mutants was calculated by dividing the number of unique sequences detected by the number of possible sequences in each class (4*C*(*L*, *d*)3^*d*^ where *C* is the combination operator to select *d* elements from a set of size *L*, where *L* = 21 is the length of the variable region, and the factor 4 is the number of families in the pool).

### Point estimation of model parameters

Model parameters *k* and *A* for each sequence were estimated using least-squares fitting on reacted fractions with different initial BYO concentrations. Least-squares fittings were performed using the ‘optimize.curve_fit’ function of the ‘SciPy’ package in python with “trust region reflective” (trf) method. The initial values of *k* and *A* were sampled from uniform distribution between 0 and 1. The bounds [0, 1] were applied on *A* values and [0, +∞) on *k* values. The tolerances for optimization termination (ftol, xtol, gtol) were kept as default (10^−8^). Optimal *k*, *A* determined from all sample points for a sequence were reported as point estimates.

### Simulated count data based on the experimental pool

To best resemble the experimental pool and conditions, we simulate the *k*-Seq pool dataset from experimentally measured values. We sampled *M* = 10^6^ sequences with parameters (*p*_*i*0_, *k*_*i*_, *A*_*i*_) estimated in the *k*-Seq experiment on the mixed variant pool, where *p*_*i*0_ is the relative abundance for sequence *i* in the unreacted pool (normalized to 1 after sampling) and *k*_*i*_, *A*_*i*_ are point estimates from fitting. We used the pseudo-first order rate equation (equation 1) to calculate the reacted fraction *f*_*ij*_ for each sequence with initial BYO concentrations from the extended substrate range and triplicate samples. The relative abundance in the simulated reacted samples was 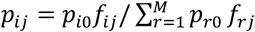. We then used the multinomial distribution Multinomial(*N*_*j*_, *p*_1*j*_, *p*_2*j*_, …, *p*_*Mj*_) (where *N*_*j*_ = 40*M*, yielding a similar mean count per sequence to that observed in the experimental pool), to model the process of sequencing by sampling *N*_*j*_ reads for each sample (unreacted and reacted). To simulate total RNA recovered, we sampled from Normal(μ_*j*_, 0.15μ_*j*_), where 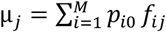 is the total RNA amount reacted in the mixed pool reaction and 0.15 is the relative error. The value of the relative error was based on the standard deviation calculated from quantification using the spike-in or by direct RNA amount quantification (Figure S2).

### Uncertainty estimation using bootstrapping

The uncertainty of estimation was assessed using bootstrap sampling of the relative residuals. Let *f*_*ij*_ be the reacted fraction for sequence *i* in reacted sample *j*, and 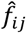 be the fitted value from point estimation. For each sequence, we calculated the relative residual as 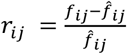. Each bootstrapping process resampled the relative residuals for sequence *i* (with replacement) to the same sample size 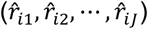, then applied the resampled relative residuals to 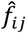 with proper scaling (i.e., 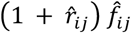 as bootstrapped data points. Least-squares fitting was performed on each set of bootstrapped data points for which *k*, *A*, and *kA* values were recorded. Sample mean, standard deviation (s.d.), median, and estimated 95% confidence interval (CI-95, as mean ± 1.96 s.d. or [2.5-percentile, 97.5-percentile)] on *k*, *A* and *kA* were calculated from bootstrapped results for each sequence.

We performed least-squares fitting on data from the mixed pool of variants (reported in this paper), data from the previously published selection (2), the simulated reacted fraction dataset (reported in this paper), and the simulated pool dataset (reported in this paper). Bootstrapping was performed for 100 re-samples for each sequence for uncertainty estimation. To compare the performance of bootstrapping, we also applied the triplicates method, used previously (2), to the simulated pool dataset, with each replicate in a BYO concentration assigned to one of three series. Each of the simulated triplicate series was fitted separately to calculate the standard deviation in estimating *k*, *A* and *kA* for each sequence.

## RESULTS

### Model identifiability depends on kinetic coefficients, experimental conditions, and measurement error

We used the simulated reacted fraction dataset to evaluate the effect of kinetic coefficients, measurement error, and experimental conditions on model identifiability for pseudo-first order kinetics, specifically if k and *A* can be separately estimated. We first evaluated the model identifiability qualitatively, for sequences selected from 6 regions over the parameter space of log_10_(*k*) from −1 to 3, log_10_(*A*) from −2 to 0, and *kA* > 0.1 min^−1^M^−1^ (Figure S3-S9). In each region, sequences fitted in repeated fitting or bootstrapping were sampled for visual evaluation of the separability of *k* and *A*, under various measurement error rates (*∊*). As summarized in Table S2, sequences with higher *k*, *A* values and lower *∊* are more likely to be separable.

To quantify separability for individual sequences, we calculated three metrics: Δ*A* or the range of *A* across repeated fittings (no resampling); *σ*_*A*_, or the standard deviation of *A* from bootstrapping samples; and *γ*, a measure of how noisy the separate estimation of *k* and *A* is compared to estimating the combined parameter *kA*. Δ*A* was able to identify sequences with numerically unstable fitting results which have small *k*, *A* values, but was not able to identify sequences whose fitting optima were sensitive to noise in the data. Almost all the sequences from the 6 selected regions each converged to a uniform optimum in repeated fitting, and the convergence was insensitive to the level of noise. In contrast, bootstrapping results account for noise in the data, and the optima from fitting subsampled data points did not always converge, providing more comprehensive separability information than convergence of multiple fittings. By comparing the distribution of each metric for sequences in the selected regions, both *σ*_*A*_ and *γ* reflected the trend of model identifiability observed by examining individual curves: higher metric value corresponded to less separable parameters of a sequence (Figure S10). In practice, we found *σ*_*A*_ and *γ* aligned equally well with human intuitions (Figure S11) and with each other (Figure S12) in evaluating model identifiability from variant pool *k*-Seq data.

Using metric *γ*, we further assessed the effects of experimental conditions and measurement error on model identifiability (Figure 2). Model identifiability depends on the true *k* and *A* values, choice of substrate concentrations, and the level of measurement error. Controlling the experimental design and measurement error, *k* and *A* were more separable for sequences with higher *k* and *A* value. Comparing the sequences for a given *kA*, model identifiability appears to be more dependent on *k* than *A*, especially for cases with lower measurement error (e.g., relative error ≤ 0.5). To assess the effect of experimental conditions, we compared the case of adding one more replicate to each reaction (4 vs. 3) to adding a higher concentration of BYO (1250 μM) in triplicate. Despite having more samples, adding another replicate did not change the region of sequences with identifiable models. However, adding a higher concentration of substrate shifted the boundary of separability on *kA* values by a factor of ~10, effectively increasing the dynamic range of the *k*-Seq assay (Figure 2A). Additionally, the difficulty of separating *k* and *A* increased when the measurements were noisier (Figure 2B). Sequences with separable parameters at low measurement error (e.g., *kA* ~ 10 min^−1^M^−1^ and relative error < 0.2) became non-separable when the measurement error was large (e.g., relative error = 1.0).

**Figure 2.**
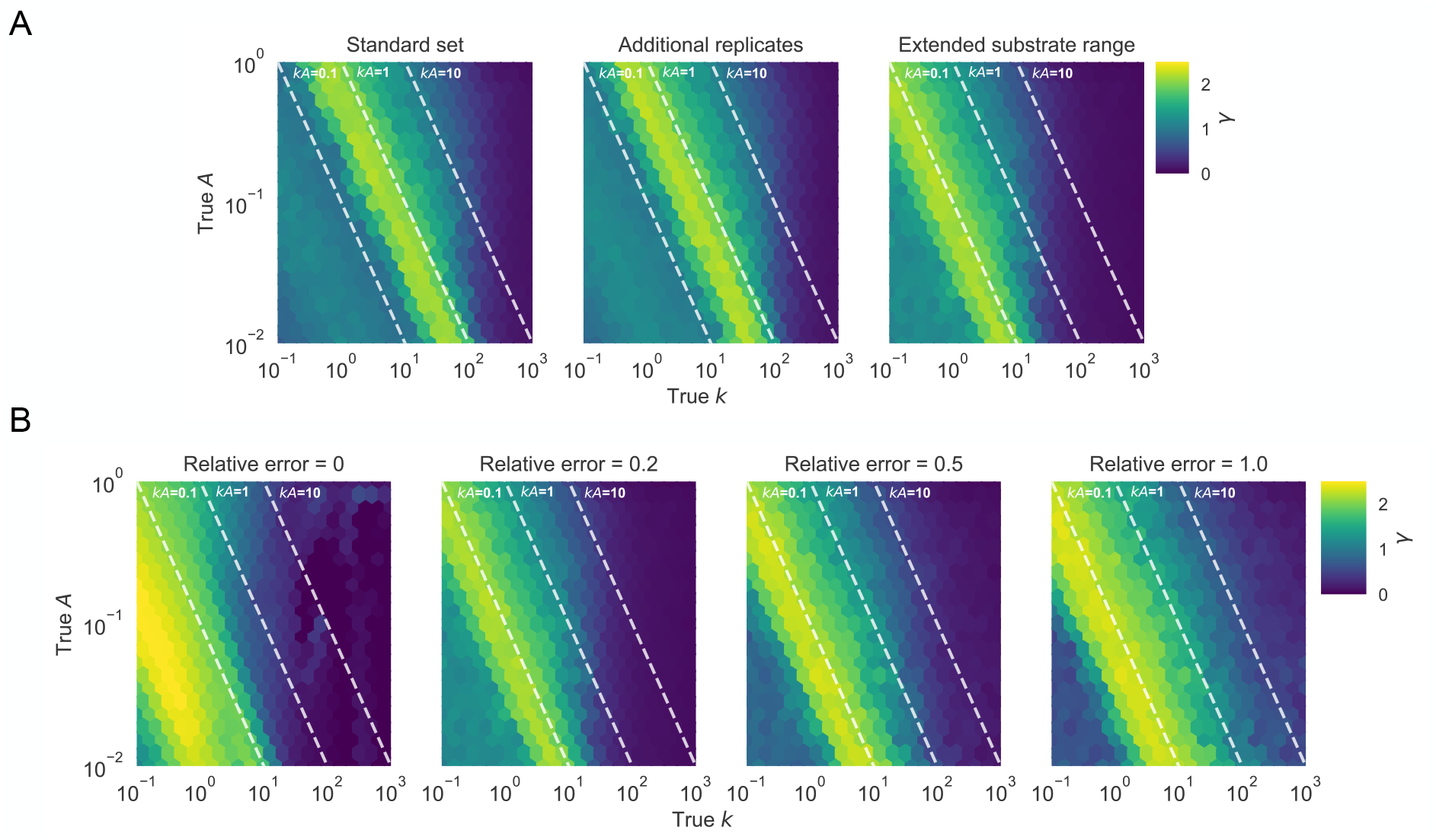
Effect of experimental factors on model identifiability to separately estimate *k* and *A*. Identifiability was evaluated using metric *γ*, based on the simulated effects of (A) choice of BYO samples (with relative error = 0.2) and (B) relative error (using the BYO series of the extended substrate range). Reacted fractions for 10,201 (101^2^) simulated sequences with true *k*, *A* in the parameter space shown in the figure were fit to the pseudo-first order model, and *γ* values for each sequence were calculated from 100 bootstrapped samples. Higher values of *γ* indicate that *k* and *A* are less separable. (A) Choosing a wider range of BYO concentration is more effective in improving the region of identifiable data compared to adding more replicates of the same BYO concentrations. (B) With higher measurement error, *k* and *A* become increasingly difficult to estimate separately.

While the ability to estimate *k* and *A* separately is of general interest for kinetic measurements, we found they could not be estimated separately for most of sequences within a Hamming distance of 2 to the family centers in the variant pool (Figure S11). Thus, for the purpose of analyzing accuracy and uncertainty for *k*-Seq over a wide range of activities (analysis below), we focus on the estimation of the combined parameter *kA*.

### *k*-Seq of ribozyme mutants: data pre-processing

We conducted the *k*-Seq experiment on a multiplexed sample containing mixed pools of variants of ribozymes S-1A.1-a, S-1B.1-a, S-2.1-a, and S-3.1-a, using the expanded experimental conditions evaluated above (2 to 1250 μM substrate). A known amount of an unrelated RNA sequence (the ‘spike-in’ sequence) was added to each reaction to aid absolute quantitation. After demultiplexing the reads, we obtained 39,151,684 paired-end reads in the unreacted sample and a mean of 13,057,929 (s.d. = 4,359,249) paired-end reads in reacted samples (Figure S13A). Around 90 - 92% of the reads were successfully joined in each sample (Figure S13C). Dereplication, removal of reads with length not equal to 21, and removal of the spike-in sequence reads yielded a count table of the number of reads for each unique sequence detected in each sample. On average 87.9% (s.d. = 1.1%) of total reads were preserved in the samples.

In principle, in order to calculate the reacted fraction with at least one non-zero value for fitting, a sequence must be detected in the unreacted sample (denominator) and in at least one of the reacted samples (numerator). Using this initial criterion, 764,756 valid unique sequences were considered to be analyzable for least-squares fitting to a pseudo-first order kinetic model, which comprised 77.9% of total reads in the unreacted sample and an average of 87.7% (s.d. = 0.6 %) of total reads among reacted samples (Figure S13BC).

### Absolute quantitation of sequence concentration

While the relative abundance of each sequence in a particular reaction sample can be calculated by dividing read counts by the total reads in each sample, calculation of the reacted fraction of each sequence requires comparing the absolute quantity of each sequence in each sample to the quantity of that sequence in the unreacted sample. This can be done by measuring the absolute RNA quantity in each sample. We compared two methods: 1) spiking in a sequence at a known concentration into each sample, providing a conversion between the number of reads and absolute concentration in each sample; or 2) measurement of the total absolute RNA concentration of each sample by Qubit or qPCR. Sample quantitation by both methods agreed well with each other (Figure 3), with both having comparable standard deviation among triplicates (Figure S2). Further analysis was done based on the second method for quantitation, since the first method is disadvantageous in reducing the HTS reads available for other sequences and requiring additional bioinformatic steps.

**Figure 3.**
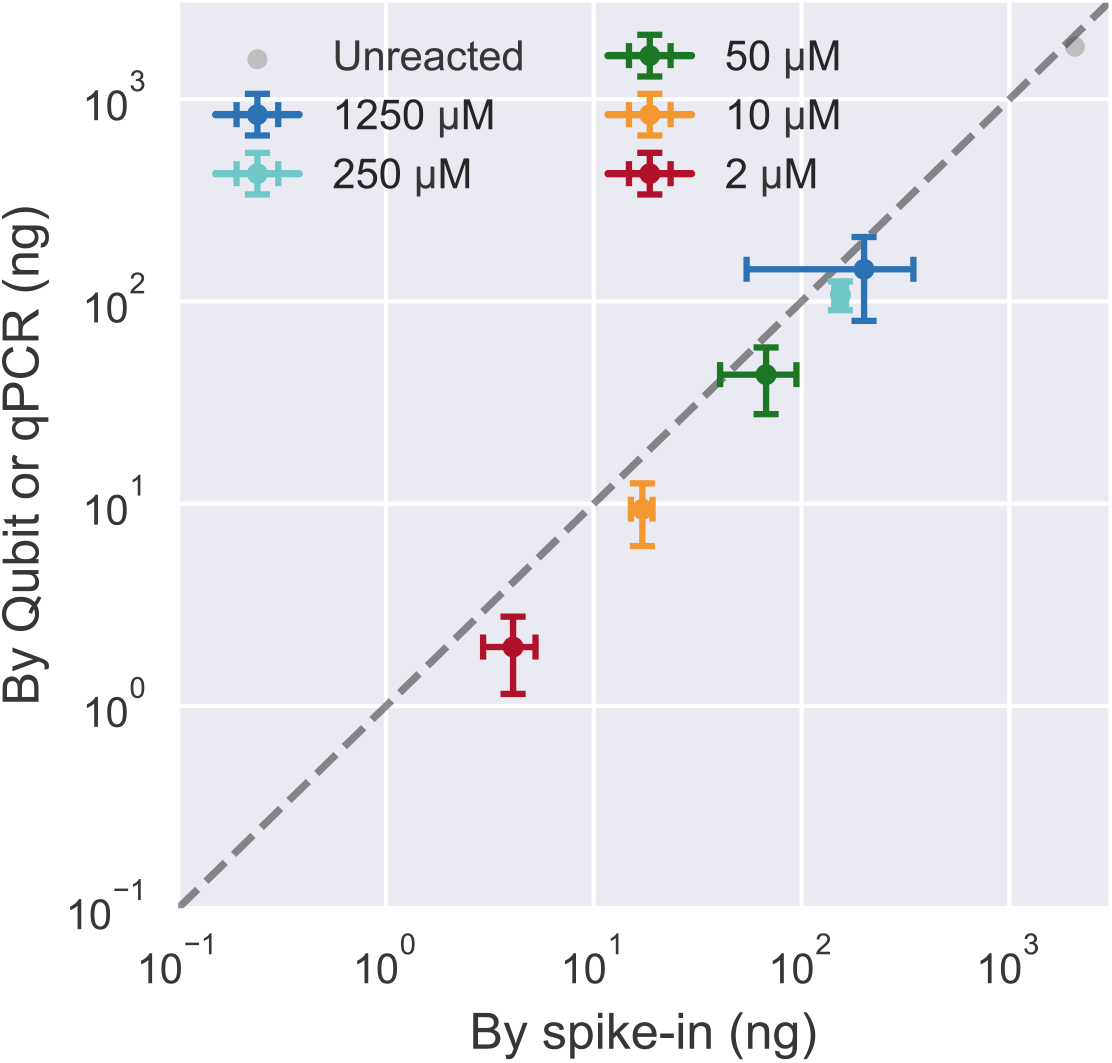
Comparison of RNA quantitation methods for *k*-Seq. Total RNA amount quantified for samples incubated with different BYO concentrations, determined by spike-in method vs. direct quantification using Qubit or qPCR, correlates well (Pearson’s *r* = 0.999, p-value = 4.78 × 10^−21^) and with comparable standard deviation (Figure S2). Error bars show standard deviations calculated from triplicates for reacted samples.

### Distribution of ribozyme mutants in the variant pool

For each chosen ribozyme (S-1A.1-a, S-1B.1-a, S-2.1-a, S-3.1-a), a variant pool was chemically synthesized such that each position had the wild-type identity with 91% probability, with non-wild-type residues being equally probable (i.e., 3% each). In theory, each variant pool should contain ~14% wild-type (*d* = 0), ~0.45% of each single mutant (*d* = 1), and ~0.015% of each double mutant (*d* = 2), or a ratio of ~0.033 for the abundance of a *d*-th order mutant to a (*d-1*)-th order mutant (Text S1, S2). The wild-type probability was selected to maximize the relative abundance of double mutants (Figure S14). We sequenced the unreacted pool and categorized each sequence read according to ribozyme families (1A.1, 1B.1, 2.1, 3.1) and the Hamming distance to the family center. Sequencing results confirmed that the mixed variant pools followed the design (Figure 4A). The variant pools contained at least ~1,000 reads per sequence for *d* ≤ 2 (up to double mutants, see Figure 4A and Table 1), and a mean of 39.7 reads per sequence for *d* = 3. Thus, the unreacted pool showed good coverage of analyzable sequences for *d* ≤ 3 (Table 1).

**Figure 4.**
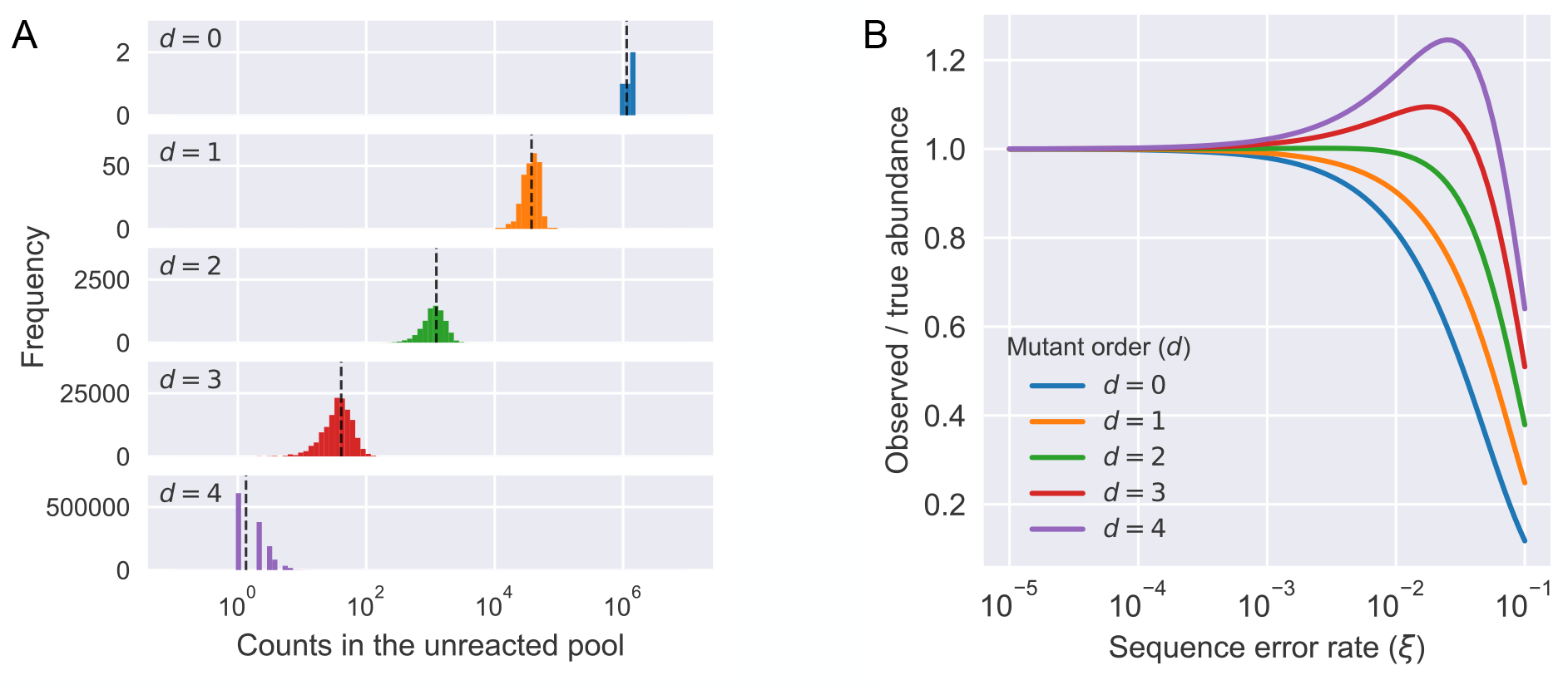
Distribution of mutants in the pool and the effect of sequencing error. (A) Relative abundance (counts) of sequences in the unreacted pool (four ribozyme families, total number of reads = 32,931,917), categorized by Hamming distance to its nearest family center. Observed abundance of different classes was similar to the expected number of counts (black dashed line). (B) The effect of different levels of sequencing error (*ξ*) to the expected observed abundance as the ratio to the true abundance for mutants with different orders (*d*) in a variant pool with 9% mutation rate. Due to the mixed effects of losing counts from being misidentified to a neighboring sequence and gaining counts from the misidentification of a neighboring sequence, the observed abundance for a sequence would either decrease (*d* = 0, 1) or first increase then decrease (*d* = 2, 3, 4) as the sequencing error increases. See Text S2 for calculation details.

**Table 1:** Coverage of local sequence space in the variant pool containing four ribozyme families. N/A = not applicable. The calculation of expected counts in the unreacted pool does not include effects of sequencing error.

| Order of mutants (*d*) | # of unique sequences | # of analyzable sequences | Fraction of possible unique sequences that were analyzable | Mean counts in the unreacted pool (std. dev.) | Expected counts in the unreacted pool |
| --- | --- | --- | --- | --- | --- |
| 0 | 4 | 4 | 1.000 | 1,243,500 (151,356) | 1,136,125 |
| 1 | 252 | 252 | 1.000 | 37,599.9 (10,607.1) | 37,455 |
| 2 | 7,560 | 7,560 | 1.000 | 1,198.3 (454.8) | 1,234 |
| 3 | 143,640 | 143,482 | 0.999 | 39.7 (18.9) | 40.7 |
| 4 | 1,939,140 | 590,115 | 0.304 | 2.1 (1.3) | 1.34 |
| ≥ 5 | N/A | 23,343 | N/A | 1.1 (1.3) | N/A |

Sequencing errors of a common sequence can spuriously inflate the apparent counts of a related sequence, a particularly acute problem if the pool is uneven and a small number of sequences (e.g., wild-type sequences) are very highly represented. Consequently, sequencing errors could potentially confound *k*-Seq results and lead to incorrect estimation of the kinetic coefficients. This may be a particular problem for less abundant or less active sequences that are closely related to more abundant and active sequences (e.g., resulting in estimation of parameters being biased toward those of the abundant sequence). There are two distinct effects related to this problem. First, the number of reads observed for a given sequence is lower than the true number, due to erroneous reads that are assigned to other sequences. Second, the number of reads observed for a sequence is inflated by the contribution of erroneous reads arising from related sequences. The combination of the two effects could change the observed sequence abundances substantially at high sequencing error rate (Figure 4B). In our variant pool, containing a 9% mutation rate, a sequencing error rate of 1% could cause more than 10% of reads for a mutant (*d* ≥ 1) to be the result of sequencing error from its neighbors (Figure S15). On the other hand, the most abundant sequences in the doped pool, wild-type sequences (*d* = 0), were least affected by the sequences from its neighbors. This problem can be mitigated by decreasing the error rate. If the sequence length is small enough to be covered by paired-end sequencing, requiring absolute matching of the overlapped region between the paired-end reads of a single sequence during joining should result in compounded fidelity (e.g., from an error frequency of 1% to 0.01%). With a sequencing error rate of 0.01%, the fraction of spurious reads from neighboring sequences was reduced to < 0.5 % for up to quadruple mutants (*d* ≤ 4) without significant loss of reads during joining (Figure S13B).

### Accuracy of *k*-Seq estimation of model parameters and uncertainty: simulated pool count data set

To evaluate the accuracy of *k*-Seq for estimating kinetic model parameters, sequences with read counts in unreacted and reacted pools were simulated using parameters (*k* and *A*) estimated from the variant pool *k*-Seq experiment as the ‘ground truth’. The data set was constructed to simulate an experiment in which ribozymes were reacted with the extended BYO concentration series in triplicate. These data were fitted according to the pseudo-first order kinetic model to estimate *k* and *A*. We expected that sequences having a low number of counts in the data set would show reduced estimation accuracy. To characterize this effect, we plotted the ratio of the point estimate of *kA* to the true *kA* against the average number of counts across the simulated samples (Figure 5A). We found that the relative error in estimation for sequences with high mean counts (> 1,000) was <10% and < 2-fold for a mean count around 100. However, the error increased substantially as the mean count decreased below 100. Thus, sequences with low mean counts (especially <100), either from low abundance in the input pool or low abundance in the reacted pool (due to low activity), were susceptible to high error in estimation. Meanwhile, very high mean counts (e.g., > 10,000) would not substantially benefit the measurement, as other sources of experimental error would likely be greater (2, 15). Thus, the results indicate that >1,000 mean count would be favorable for estimation, with >100 counts being acceptable if a 2-fold error in estimation is tolerated.

**Figure 5.**
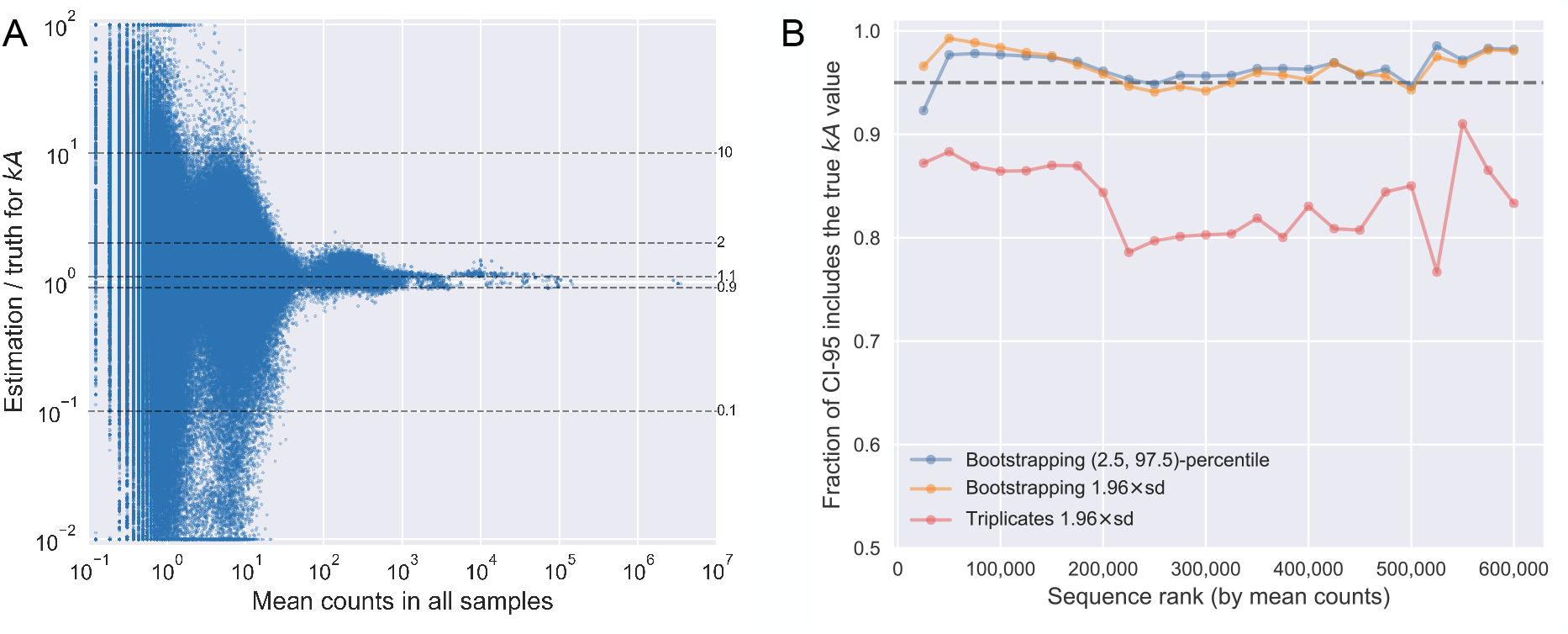
Accuracy of parameter estimation by *k*-Seq. (A) Dependence of accuracy (ratio of estimated *kA* to true *kA*) on mean counts across all simulated samples (including the unreacted pool sample). The dashed lines correspond to ratios as labeled. Ratios above 100-fold or below 0.01-fold are shown at the borders of the plot. (B) Fraction of sequences for which the CI-95, estimated using bootstrapping or using triplicates, includes the true *kA* values, for sequences with different mean counts across all samples. Sequences were ranked by mean counts (from highest to lowest) and binned in sets of 25,000 sequences. Each data point indicates the fraction of CI-95 that includes the true values in each bin.

While the above analysis demonstrated the accuracy of point estimation, a reliable quantification of uncertainty is required to assess the precision in estimating from real data when the ground truth is unknown. We therefore explored the accuracy of uncertainty quantification using bootstrapping. Bootstrap resampling (n = 100) was used to estimate the 95% confidence intervals (CI-95) in two ways: first, using mean and standard deviation of estimated *kA* (mean ± 1.96 s.d., assuming a normal distribution), and second, using the 2.5-percentile to 97.5-percentile confidence intervals (normal distribution not assumed), for sequences in the simulated pool dataset that were analyzable (602,246 sequences in total). A sensible evaluation of CI-95 estimation is the fraction of sequences with true *kA* value included in the estimated CI-95. If the estimation were correct, the CI-95 would include the true value for roughly 95% of sequences. We found bootstrapping gave an accurate CI-95 estimation by either method. 96.5% of sequences included the true *kA* within the estimated CI-95 from 2.5-to-97.5-percentiles, and 96.4% did so when estimating CI-95 from the mean and standard deviation. Of note, the fractions of sequences with their true *kA* included in the CI-95 were relatively consistent regardless of their mean counts (Figure 5B) or true *kA* values (Figure S16), indicating that these methods could be used broadly to quantify uncertainty for sequences across different abundance or activity values. For comparison, we also examined uncertainty estimation using the mean and standard deviation estimated from triplicates (mean ± 1.96 s.d.). In our simulated pool dataset, 83.5% of sequences had the true *kA* value included in the CI-95 estimated from triplicates, indicating that uncertainty is underestimated for a substantial fraction of sequences (i.e., over-confident in estimated values) (Figure 5B).

### Precision of *k*-Seq estimation: experimental data set

The precision of *k*-Seq measurement for data from the variant pool experiment was evaluated in two ways. First, given the reasonable accuracy obtained by the bootstrapping procedure, we calculated the fold-range (97.5-percentile divided by 2.5-percentile) estimated from bootstrapping (n = 100). While there was a slight tendency for sequences with higher *kA* to have higher estimation precision (lower fold-range; Figure S17) in each order of mutants, the precision was more evidently dependent on the mean counts value for sequences, both within and across groups (Figure 6A). All wild-type sequences and most single and double mutants had CI-95 spanning less than one order of magnitude (fold-range < 10). Triple mutants had generally lower precision, consistent with their lower counts.

**Figure 6:**
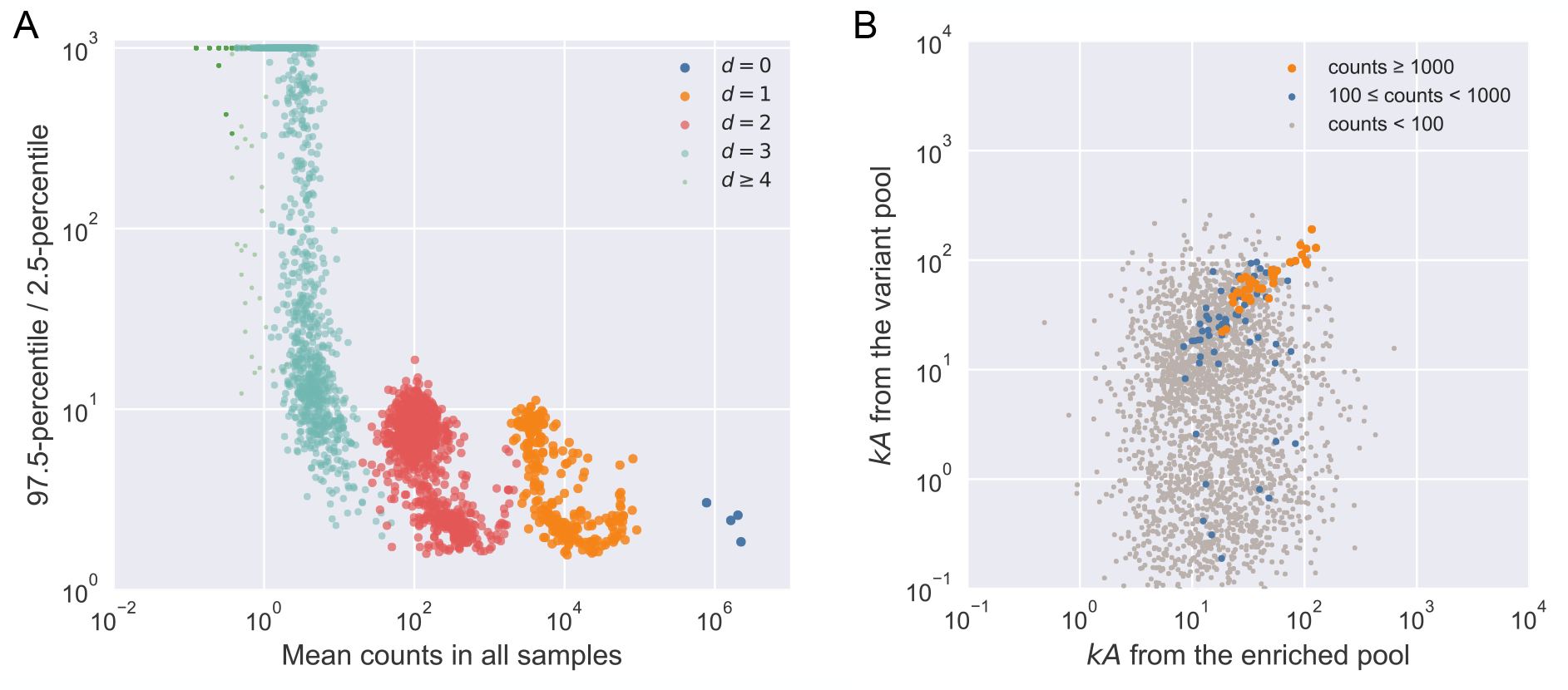
Precision of estimation by *k*-Seq. (A) Fold-range (97.5-percentile / 2.5-percentile) of *kA* estimation depended on the mean counts. Increasing mean counts increases precision, as shown by the relationship of fold-range with mean counts across different orders of mutants. For *d* ≥ 2, only 1000 sequences were randomly selected for visualization. (B) Alignment between estimated *kA* from two independently conducted experiments (experiment from (2), and the *k*-Seq experiment reported here). Only sequences with 2.5-percentile higher than baseline catalytic coefficient (*kA* = 0.124 min^−1^M^−1^, reported in (2)) were included. Each point represents a sequence whose color reflects the minimum of mean counts (between two experiments).

Precision as measured above represents variation among replicates done in the same experimental batch but does not include variation between different *k*-Seq experiments. To understand the precision of estimates from independently designed and separately executed *k*-Seq experiments, we compared the results from the variant pool *k*-Seq reported here to a previously reported *k*-Seq assay from a selection pool (2). 2513 unique sequences were found for which the 2.5-percentile for *kA*, estimated from bootstrapping, was greater than the baseline *kA* of 0.124 min^−1^M^−1^ (measured in (2)) in both experimental data sets. Point estimates of *kA* for these sequences from the two experiments were compared to each other (Figure 6B). For sequences with sufficient counts (e.g., mean counts of at least 1000 in both experiments, corresponding to 39 sequences), the results from the two experiments were well correlated (Pearson’s *r* = 0.896, p-value = 1.20 × 10^−14^; Spearman’s ρ = 0.864, p-value = 1.38 × 10^−12^), indicating good repeatability of those measurements from different experiments. As expected, lowering the count threshold gave a decrease in repeatability: sequences with mean counts greater than 100 in both experiments (98 sequences) showed Pearson’s *r* = 0.710 (p-value = 2.73 × 10^−16^) and Spearman’s ρ = 0.616 (p-value = 1.53 × 10^−11^). For analyzable sequences with mean counts <100 in either experiment, weak to no correlation was found (Pearson’s *r* = −0.0223, p-value = 2.73 × 10^−1^; Spearman’s ρ = 0.0517, p-value = 1.10 × 10^−2^).

## DISCUSSION

In this work, we addressed several issues related to the rigor of inferring kinetic model parameters from *k*-Seq analysis, namely model identifiability, sequencing error, and estimation of uncertainty.

A model is not identifiable when the optimal set of parameters fitting the model cannot be uniquely determined due to noise in the data (12, 16). For the ribozymes exhibiting pseudo-first order kinetics studied here, the parameters *k* (rate constant) and *A* (maximum reaction amplitude) could not be separately estimated when the data collected did not show saturation behavior, i.e., the data fell into the initial linear region of the curve. While it would be possible to adjust the substrate concentrations to mitigate the problem for individual sequences, it is impossible in *k*-Seq experiments to apply optimal conditions for each sequence due to pool heterogeneity. We previously reported the combined parameter *kA* as the measure of chemical activity (2). However, separate estimation of *k* and *A* is still an important goal. In general, we found that higher values of *k* or *A* and lower noise level yield better separability. Of the metrics (σ_*A*_ and γ) we calculated from bootstrapping results, both showed good performance scoring the separability of *k* and *A*. These metrics allows one to semi-quantitatively assess separability, in combination with experimental intuition, to determine which results can be reasonably reported as *k* and *A* separately vs. reported as the combined activity parameter *kA*.

Using a simulated data set, we studied how an experiment might be designed to increase the number of sequences in the separable region. Extending the substrate concentration range (to 1250 μM) expands the region of separable sequences by pushing the lower bound on *k* or *A* down by roughly one order of magnitude. By contrast, adding another replicate to each substrate concentration did not substantially change the separability map. Thus, we used the extended substrate range in the present *k*-Seq experiment to provide a wider dynamic range. Even so, a substantial fraction of sequences exhibited non-separable *kA*, so in practice the choice of parameters to be reported depends on the goals of the experiment (e.g., for maximum exploitation of the data, all of *k*, *A*, and *kA* could be reported along with σ_*A*_ or γ).

Using DNA sequencing counts to quantify the abundance of sequences has two consequences that need consideration: sequencing error that mis-identifies a sequence as a related sequence, and stochastic noise in measurement for sequences associated with lower counts. The first is a particular problem when sequences are close in the sequence space (e.g., mutational analysis on a variant pool) and the pool is quite uneven. In this case, sequencing errors can contribute a non-trivial portion of reads to less abundant neighbor sequences, effectively mixing the *k*-Seq measurement for these less abundant sequences with those of their neighbors. While model-based sequencing error correction could be attempted (17, 18), this problem can also be circumvented by paired-end reads. By enforcing absolute matching during the joining process, the error rate of sequencing (e.g., approximately 1% per base) can be substantially decreased (e.g., to 0.01% per base) if the entire sequence was read in both directions. While this decreases the number of reads that passed quality control, the benefit is important as misidentification from this level of sequencing error was essentially negligible for observed abundances (Figure 4A).

Low counts had a major effect on the accuracy of estimation of kinetic coefficients. In practice, we found that the average counts for a sequence across samples (i.e., mean counts) was a better guide for estimation accuracy compared to counts in the input pool (Figure 5A, Figure S18). Accuracy was good for sequences above a certain threshold of mean counts (e.g., 100 reads per sample) but decreased markedly below this. At the same time, the benefit from large counts (e.g., 10,000 reads per sample) was marginal and other experimental factors likely contribute greater error.

Estimating uncertainty is important for *k*-Seq experiments, but replicates are likely to be limited due to the expense associated with HTS. Bootstrapping simulates virtual replicates by resampling data (in this case, the relative residuals from fitting) with replacement to its original size. The results can be used to estimate population characteristics, such as confidence intervals for estimated model parameters. Indeed, bootstrapping results reflected the true 95% uncertainty level more appropriately than the standard deviation estimated from triplicate experiments, as the latter tended to underestimate the uncertainty. As seen for the accuracy of low count sequences, precision showed a steep drop-off when mean counts dropped below 100 (Figure 6A), while large counts also did not significantly improve precision. Using bootstrapping instead of replicates also provided resampling data to calculate σ_A_ and γ for model identifiability analysis. As modern computational resources become cheaper and easier to access, bootstrapping, despite its higher computation cost, becomes more affordable. Therefore, while experimental replicates are valuable for controlling for some sources of error, we suggest that bootstrapping analysis is an excellent method for properly estimating errors and understanding model identifiability.

To maximize coverage, or the number of sequences with estimated model parameters having acceptable accuracy and precision, it is desirable to maximize the number of sequences satisfying a minimal count requirement without spending excessive sequencing resources on abundant sequences. In the present experiment, we had approximately full coverage for single and double mutants in each family, for which the measurement precision may be considered reasonable (fold-range < 10; mean counts > 10) (Figure 6A). While HTS technology enabled the kinetic measurement for large pools with high richness (number of unique sequences), the practical coverage for *k*-Seq is affected by pool evenness, as highly uneven pools may have many sequences with insufficient counts for precise estimation. Such pools may result from enrichment after selections or from variant pool synthesis of ribozyme variants exploring many mutants of a given wild type. For enriched pools from selection experiments, the pool evenness usually decreases during the selection. For doped pools, evenness is tuned by the ratio of wild-type nucleotides at each position. In the analysis of BYO-aminoacylation ribozymes presented here, the designed variant pool was more even than the selection pool from which these ribozymes were derived (Figure S19); thus *k*-Seq analysis of selected ribozymes may be improved by designing a new pool rather than directly analyzing the selected pool itself.

Greater evenness in the samples may be achieved in the near future by emerging technologies for synthesizing uniform oligo pools with very high richness (19), as well as by improvements in sequencing technology to achieve greater total sequencing depth. With bootstrapping, data with low counts can be assessed with estimated uncertainty rather than discarded. On the other hand, systematic errors due to sequencing errors cannot be assessed by replicates or bootstrapping analysis; instead, effort would be well-spent reducing sequencing errors, and the degree to which results might be biased by the resulting error rate should be kept in mind when interpreting the data. Attention to these issues is important for fulfilling the promise of kinetic sequencing and related techniques for providing unprecedented insights into genotype-phenotype relationships.

## SUPPLEMENTARY DATA

See Supplementary Information.

## ACKNOWLEDGEMENT

Use was made of computational facilities purchased with funds from the National Science Foundation (CNS-1725797) and administered by the Center for Scientific Computing (CSC). The CSC is supported by the California NanoSystems Institute and the Materials Research Science and Engineering Center (MRSEC; NSF DMR 1720256) at UC Santa Barbara.

