## Supplementary information for "Kinetic sequencing (*k*-Seq) as a massively parallel assay for ribozyme kinetics: utility and critical parameters"

Yuning Shen[1], Abe Pressman[2], Evan Janzen[1,3] and Irene Chen[1,3,4,*]
[1] Department of Chemistry and Biochemistry, University of California, Santa Barbara, California 93106, United States
[2] Department of Chemical Engineering, University of California, Santa Barbara, California 93106, United States
[3] Program in Biomolecular Sciences and Engineering, University of California, Santa Barbara, California 93106, United States
[4] Department of Chemical and Biomolecular Engineering, Department of Chemistry and Biochemistry, University of California, Los Angeles, California 90095, United States

**Table S1**. Wild-type sequences selected from (1) for the variant pool

| # | Ribozyme | Sequence (random region) |
|---|---|---|
| 1 | S-2.1-a | ATTACCCTGGTCATCGAGTGA |
| 2 | S-1A.1-a | CTACTTCAAACAATCGGTCTG |
| 3 | S-1B.1-a | CCACACTTCAAGCAATCGGTC |
| 4 | S-3.1-a | AAGTTTGCTAATAGTCGCAAG |

**Figure S1**. Standard curve for qPCR measurement, fitted by ‘stats.linregress’ function from ‘SciPy’ Python package.

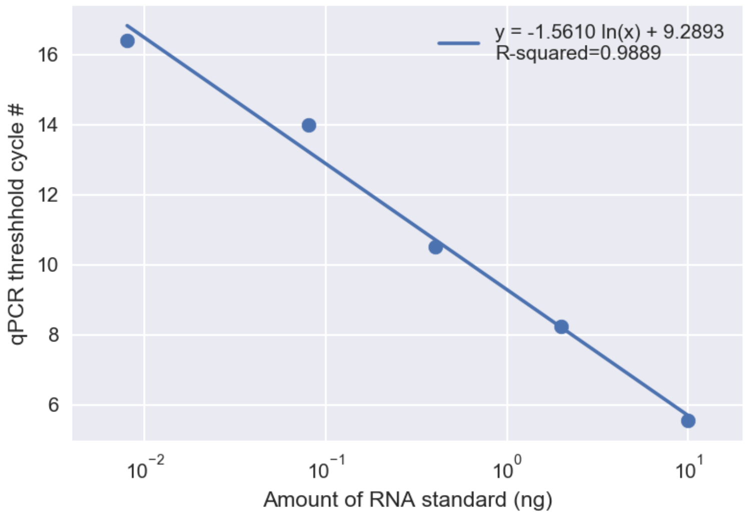

**Figure S2**. Relative standard deviation vs. mean of total RNA measured for each set of reacted sample triplicates, quantified by spike-in sequence (circles) or qPCR + Qubit (crosses). The mean relative standard deviations are similar for spike-in method (0.162) and for qPCR + Qubit method (0.176).

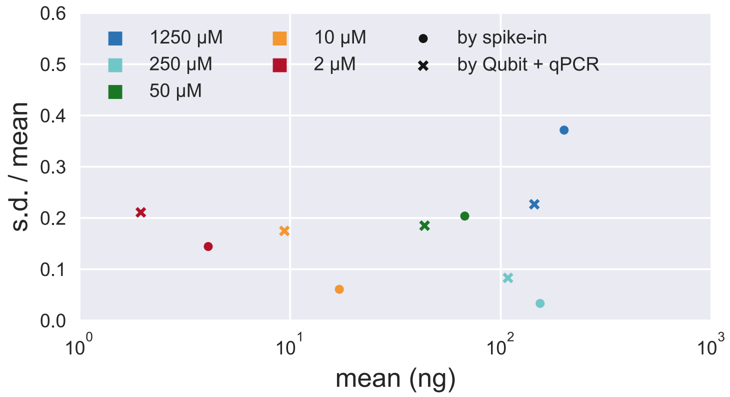

**Figure S3**. Illustration of the 6 regions selected to sample sequences and their fitting values from simulated reacted fraction dataset, with the boundary values for true $A$, $k$, and $kA$ indicated in the table (N/A = not applicable). These regions are separately analyzed in Figure S4-S9.

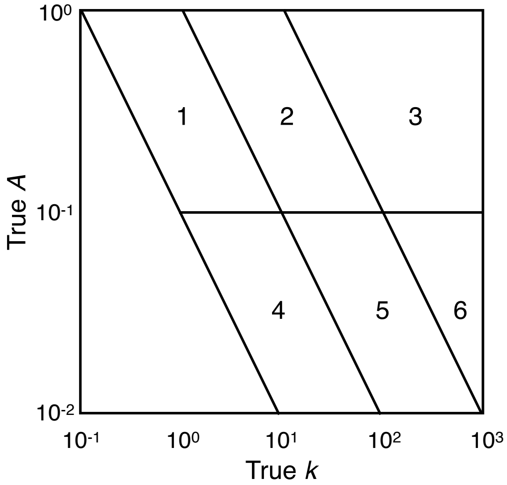


| Region | $k$ ($\mathrm{min}^{-1}\mathrm{M}^{-1}$) | $A$ | $kA$ ($\mathrm{min}^{-1}\mathrm{M}^{-1}$) |
|---|---|---|---|
| 1 | N/A | $10^{-1} < A < 1$ | $10^{-1} < kA < 1$ |
| 2 | N/A | $10^{-1} < A < 1$ | $1 < kA < 10^1$ |
| 3 | $k < 10^3$ | $10^{-1} < A < 1$ | $kA > 1$ |
| 4 | N/A | $10^{-2} < A < 10^{-1}$ | $10^{-1} < kA < 1$ |
| 5 | N/A | $10^{-2} < A < 10^{-1}$ | $1 < kA < 10^1$ |
| 6 | $k < 10^3$ | $10^{-2} < A < 10^{-1}$ | $kA > 1$ |

**Figure S4**. Selected fitting results from Region 1 ($0.1 < A < 1$, $0.1 < kA < 1\ \mathrm{min}^{-1}\mathrm{M}^{-1}$) in simulated reacted fraction dataset with different relative error. Each curve plot shows the simulated reacted fraction (in triplicates) at various initial BYO concentrations (orange crosses), fitting curves from point estimation (blue line), and fitting curves from 20 repeated fitting or 20 bootstrapped samples (grey lines); fitted $k$, $A$ values for the curves are shown in the corresponding heatmap (red crosses) under each curve plot. For visual guidance, background color of the heatmap indicates the relative values of mean squared error (normalized in each plot; blue to yellow is lower to higher error) over the parameter space given the data. The white dashed line marks $kA = 1\ \mathrm{min}^{-1}\mathrm{M}^{-1}$. An ideal fitting result would have converged fitting optima and is both numerically stable (from repeated fitting) and robust to noise (from bootstrapping). A large variance along the line of $kA = \mathrm{constant}$ indicates the model is not identifiable, i.e., $k$ and $A$ cannot be separately estimated. (react. frac. = reacted fraction.)

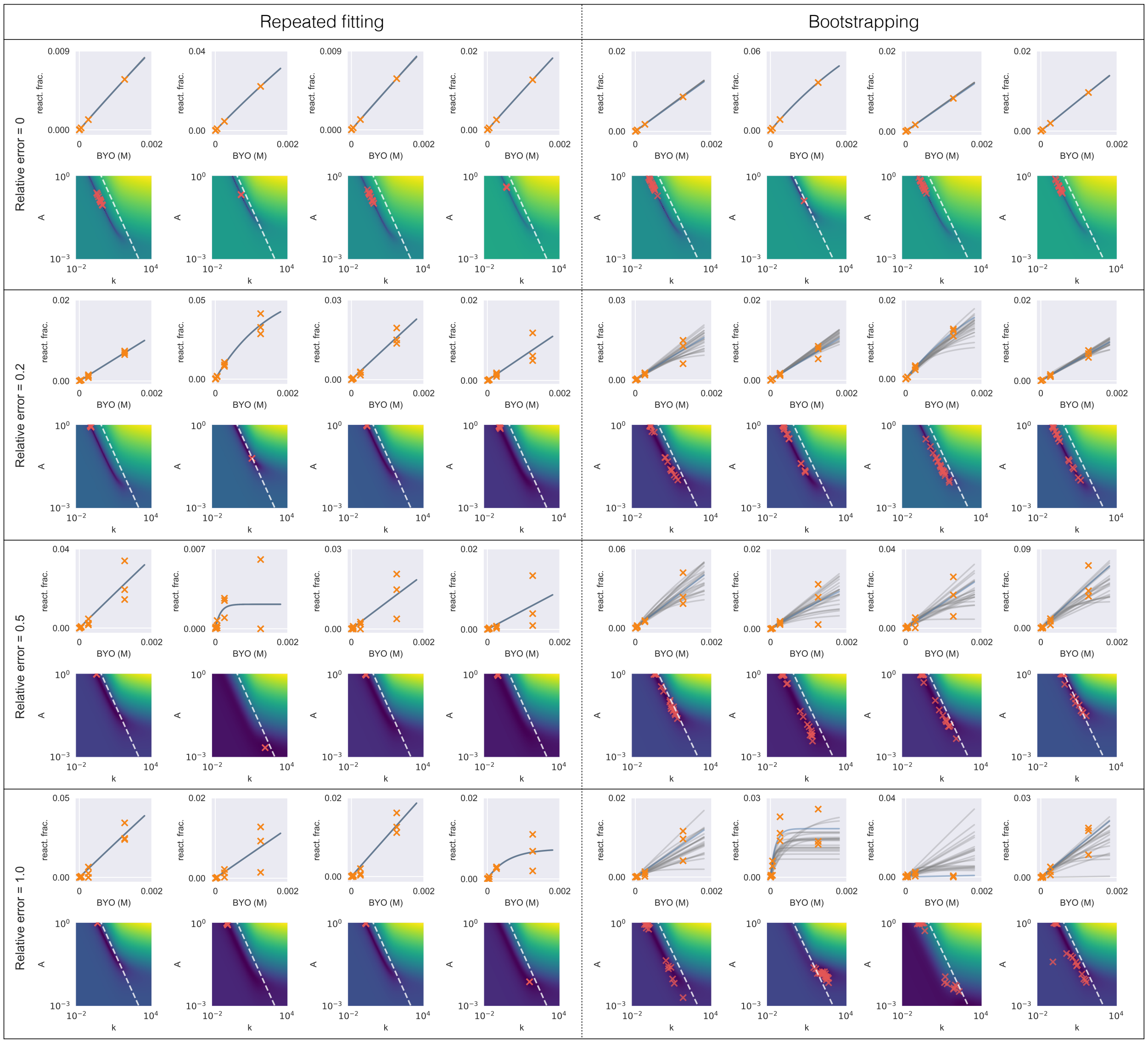

**Figure S5**. Selected fitting results from Region 2 ($0.1 < A < 1, 1 < kA < 10\ \mathrm{min}^{-1}\mathrm{M}^{-1}$) in simulated reacted fraction dataset with different relative error. See caption of Figure S4 for explanation.

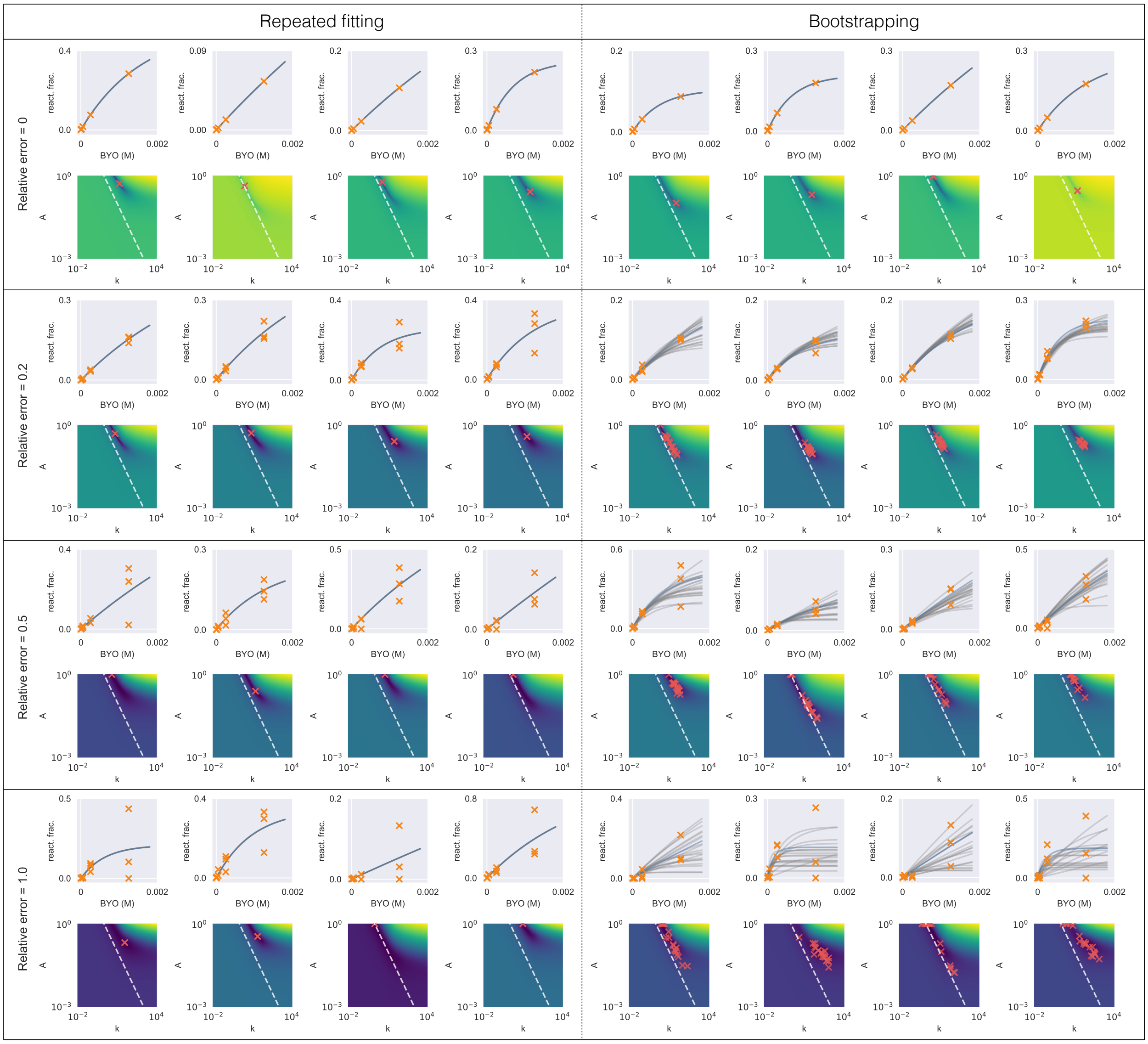

**Figure S6**. Selected fitting results from Region 3 ($0.1 < A < 1, kA > 10\ \mathrm{min}^{-1}\mathrm{M}^{-1}$) in simulated reacted fraction dataset with different relative error. See caption of Figure S4 for explanation.

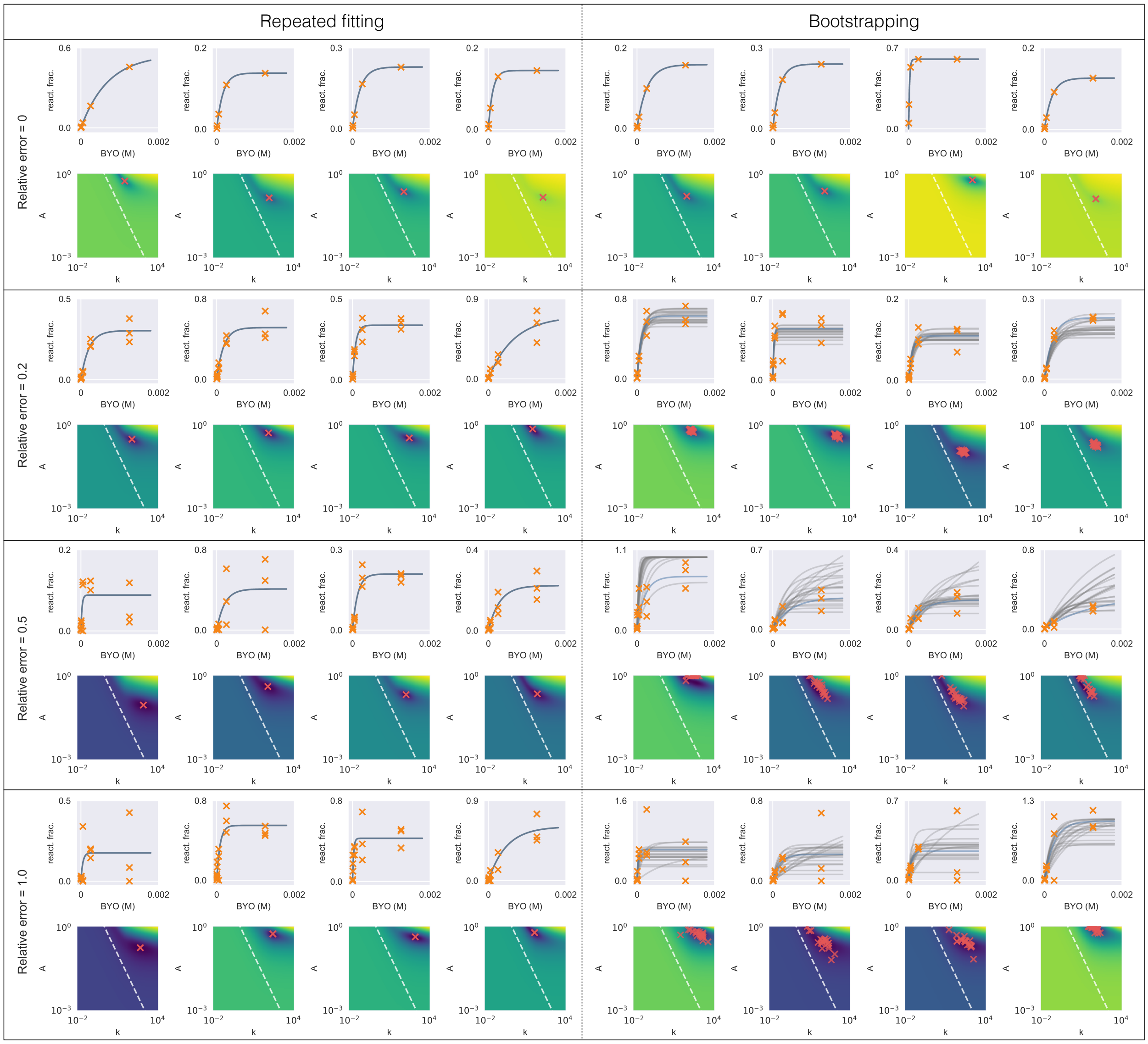

**Figure S7**. Selected fitting results from Region 4 ($A < 0.1$, $0.1 < kA < 1$ $\text{min}^{-1}\text{M}^{-1}$) in simulated reacted fraction dataset with different relative error. See caption of Figure S4 for explanation.

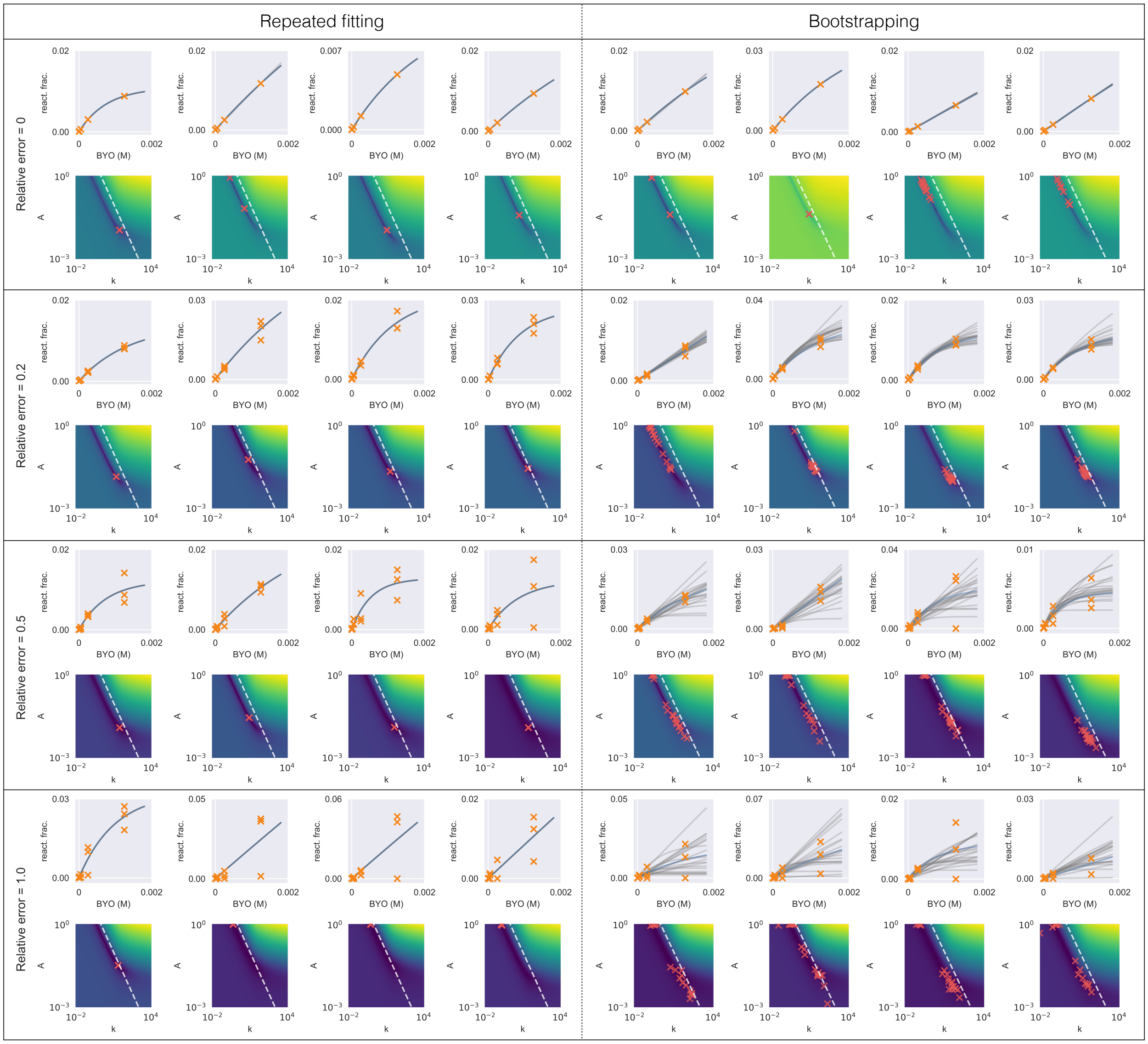

**Figure S8**. Selected fitting results from Region 5 ($A < 0.1, 1 < kA < 10 \text{ min}^{-1}\text{M}^{-1}$) in simulated reacted fraction dataset with different relative error. See caption of Figure S4 for explanation.

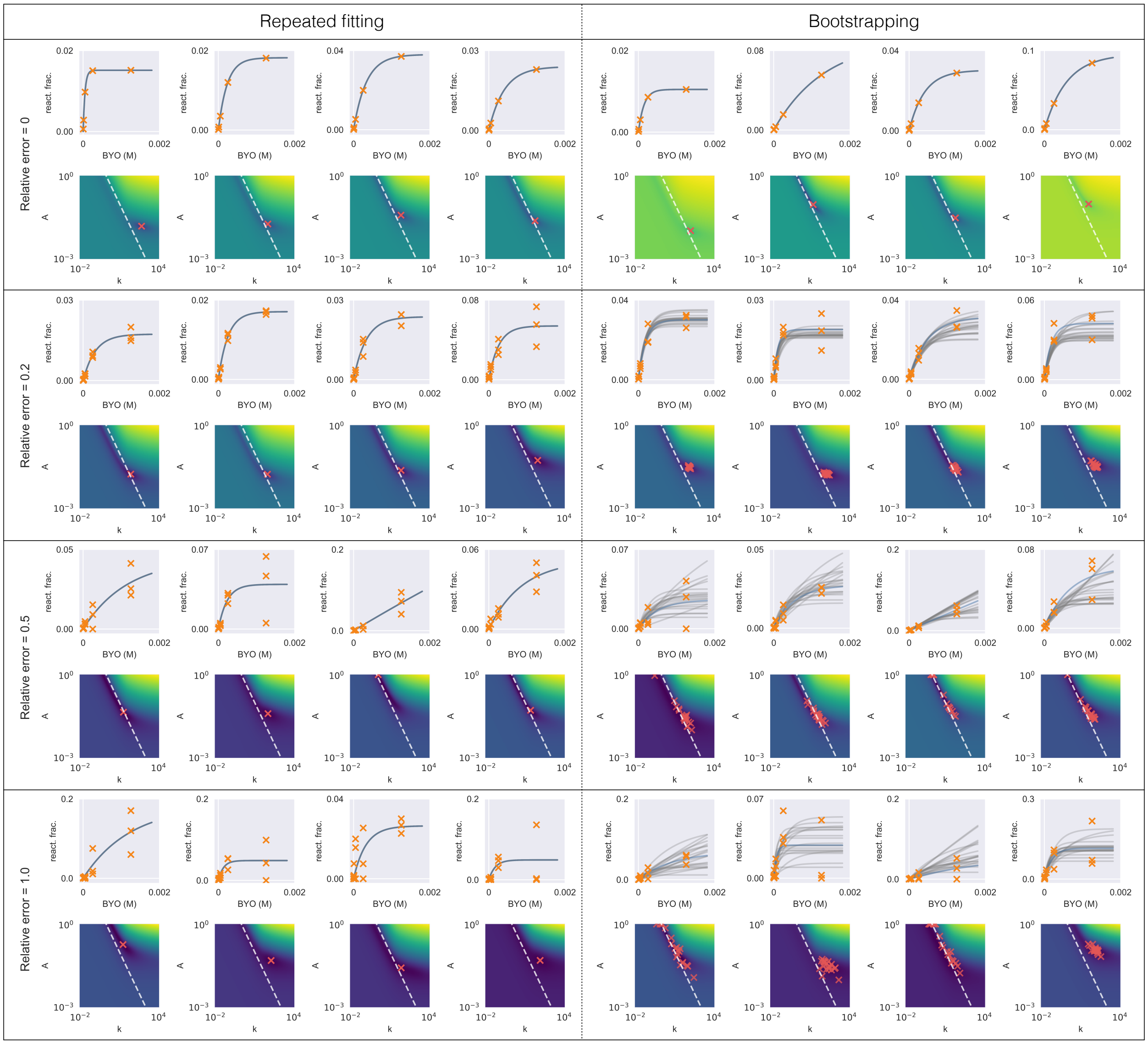

**Figure S9**. Selected fitting results from Region 6 ($A < 0.1$, $kA > 10\ \mathrm{min}^{-1}\mathrm{M}^{-1}$) in simulated reacted fraction dataset with different relative error. See caption of Figure S4 for explanation.

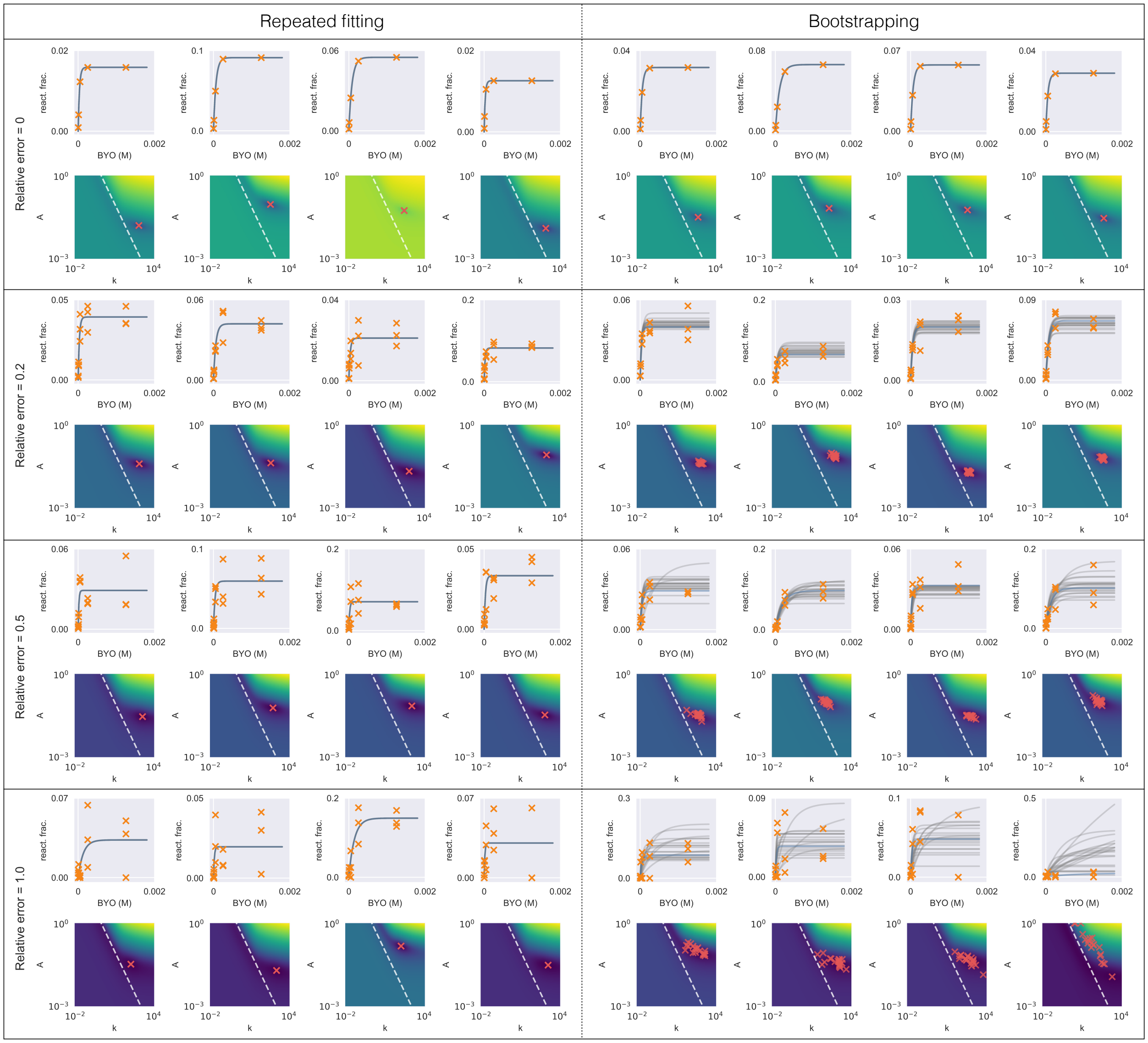

**Table S2**. Summary of visual examination of model identifiability from repeated fitting (no resampling) and bootstrapping. 'Y' indicates regions where $k$ and $A$ appear that they can be separately estimated, 'N' indicates regions where they do not appear to be separately estimable. Results from repeated fitting account for the numeric effect from different initial values and results from bootstrapping also account for the effect of sample noise.

| Method | Relative error ($\epsilon$) | Region 1 | Region 2 | Region 3 | Region 4 | Region 5 | Region 6 |
|---|---|---|---|---|---|---|---|
| Repeated fitting | 0.0 | N | Y | Y | N | Y | Y |
| | 0.2 | Y | Y | Y | Y | Y | Y |
| | 0.5 | Y | Y | Y | Y | Y | Y |
| | 1.0 | Y | Y | Y | Y | Y | Y |
| Bootstrapping | 0.0 | N | Y | Y | N | Y | Y |
| | 0.2 | N | N | Y | N | Y | Y |
| | 0.5 | N | N | N | N | N | N |
| | 1.0 | N | N | N | N | N | N |

**Figure S10**. Distribution of metric values for sequences from 6 selected regions from the simulated reacted fraction dataset, with various noise level. Histogram bars are stacked for visibility. Both metrics $\sigma_A$ and $\gamma$ captured the trend of model identifiability as summarized in Table S2. In contrast, $\Delta A$ failed to capture the difference in model identifiability between sequences with different regions and sample noise.

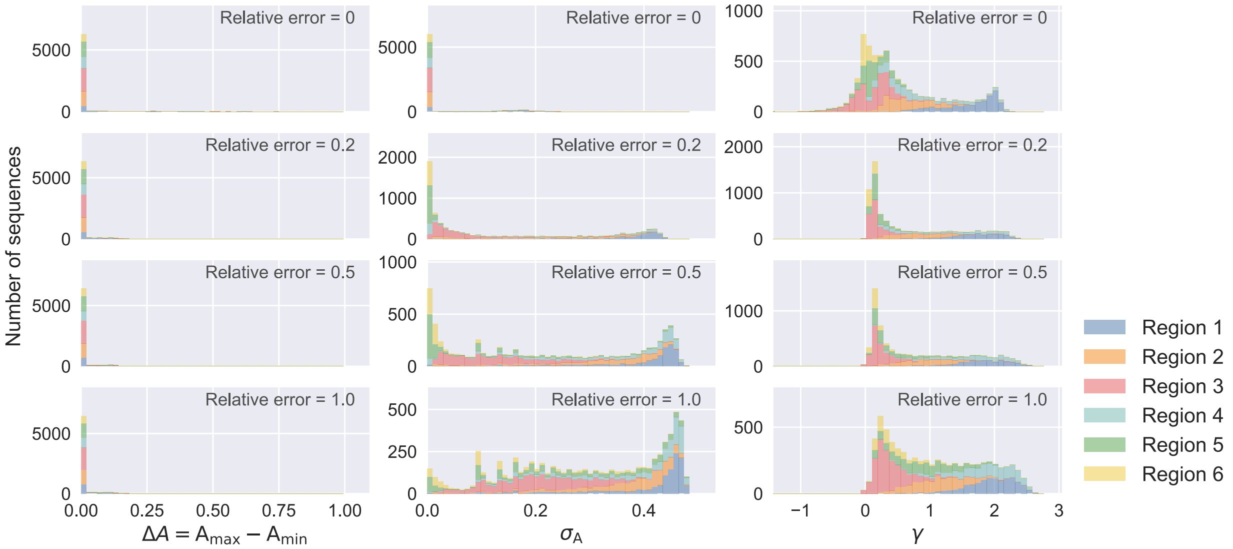

**Figure S11**. Distribution of $\gamma$ (A) and $\sigma_A$ (B) for sequences within Hamming distance of 2 to the family centers from the variant pool *k*-Seq experiment. Example fitting results are shown for sequences within each score range (labels on the left) of $\gamma$ (C) and $\sigma_A$ (D). For explanation of (C) and (D), also see caption of Figure S4. Sequences with low metric scores for both metrics showed good model identifiability while $k$ and $A$ cannot be separately estimated for those with high metric scores. $k$ and $A$ for most sequences up to double mutants could not be estimated separately.

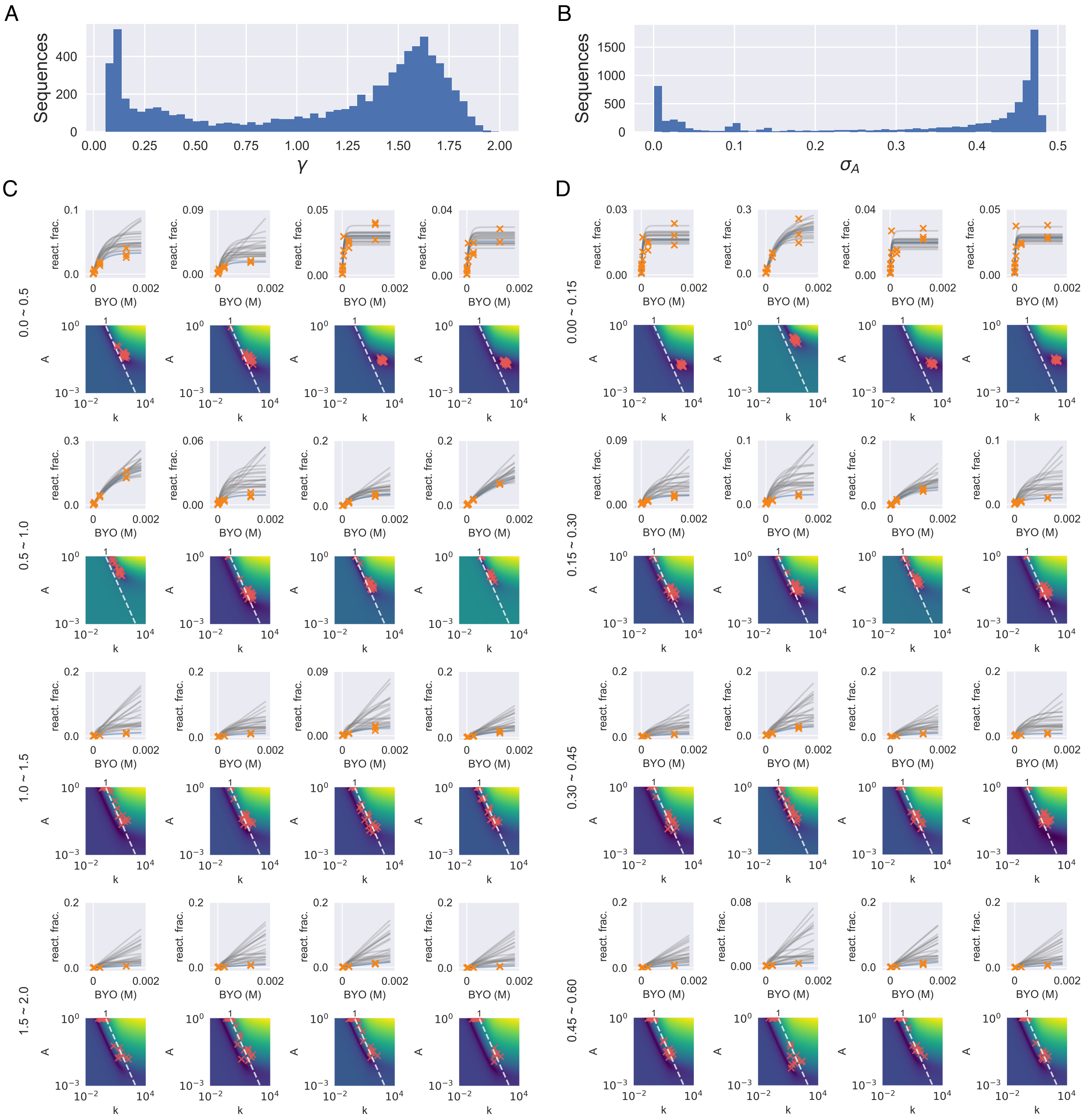

**Figure S12**. Correlation between model identifiability metrics $\gamma$ and $\sigma_A$ for analyzable sequences in the variant pool *k*-Seq experiment. Sequences within Hamming distance of 2 to the family centers ($d \leq 2$) showed good correlation between the two metrics (Spearman's ρ = 0.945, p-val = 0.000); larger variance was observed for sequences with $d > 2$ due to lower counts and noisier measurements.

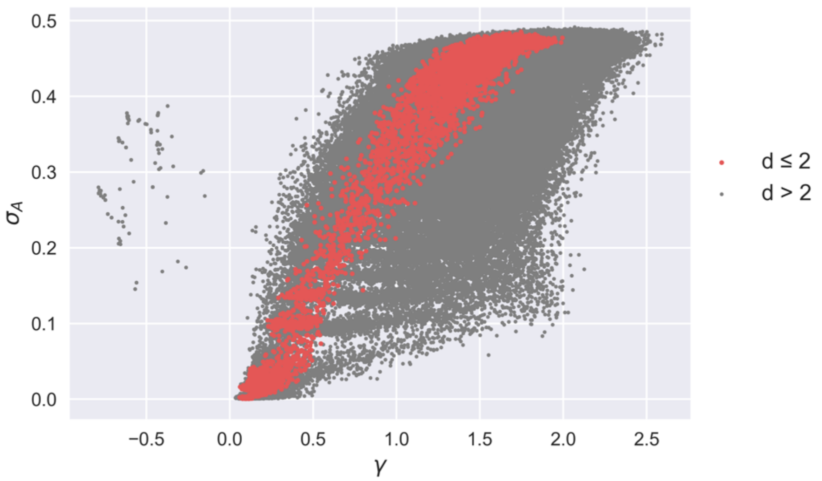

**Figure S13**. Processing of sequence reads. (A) Number of raw paired-end reads in each sample. The input sample (unreacted) had 3x of other samples for total DNA input for sequencing. (B) Number of unique sequences and (C) percent of total reads retained after paired-end reads joining, filtering (removal of spike-in sequence and sequences that are not 21 nt long), and checking for analyzability (has non-zero counts in the input pool and in at least one of the reacted samples)

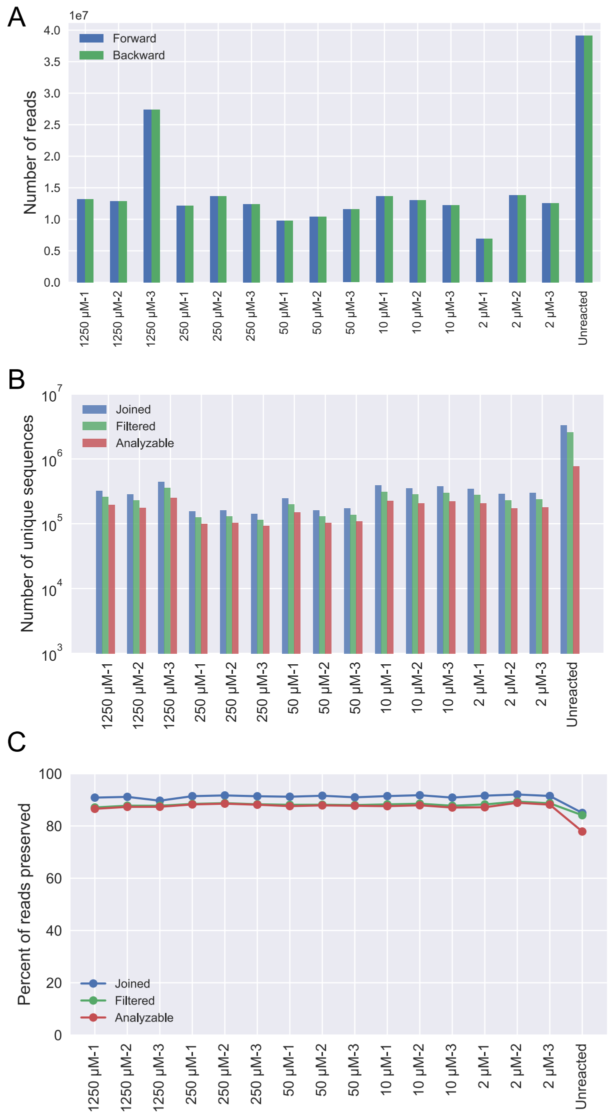

**Text S1**. Variant pool design.

We defined a sequence with *d* substitutions (Hamming distance) compared to the wild-type sequence as a *d*-th order mutant. For each partially mutated library (a single family) with randomized length $L$=21 residues and mutation rate $\eta$, the fraction of a *d*-th order mutant in the pool is

$$p_d = (1-\eta)^{L-d} \left(\frac{\eta}{3}\right)^d \tag{S1}$$

The $\eta$ maximizing the fraction of a single *d*-th order mutant satisfies $dp_d/d\eta = 0$, thus

$$\eta_\mathrm{d} = \arg\max_\eta p_d = \mathrm{d/L} \tag{S2}$$

and the maximum fraction for a *d*-th order mutant in the variant pool is

$$p_{d,\max} = \left(1-\frac{d}{L}\right)^{L-d} \left(\frac{d}{3L}\right)^d \tag{S3}$$

We used Equation S3 to determine the optimal $\eta$ maximizing the fraction of a given order of mutant in the pool. For a sample with $N$ total reads, the expected counts for a *d*-th order mutant is $p_d N$.

**Figure S14**. Expected counts for mutants in mixed variant pools of four wild types given different mutation rate (η) and $10^7$ total reads, calculated using Equation S1. For mutants with $d > 2$, the maximum number of expected counts is not sufficient for all possible *d*-th order mutant sequences to be estimated accurately (i.e., counts are < 10-100), showing a limitation of the variant pool library.

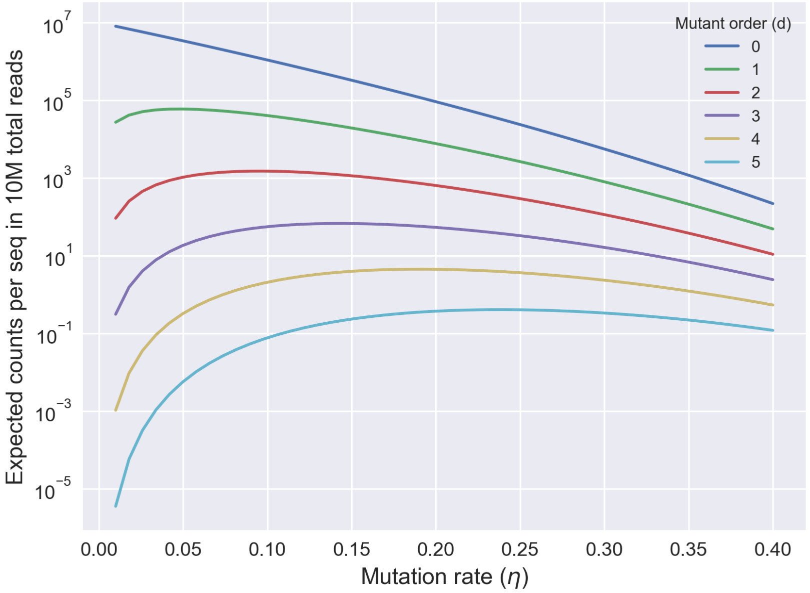

**Text S2**. Effects of sequencing error in the variant pool.

From Equation S1, the relative abundance ratio between a *d*-th mutant and a (*d+1)*-th mutant is

$$\phi = p_d/p_{d+1} = \frac{3\,(1-\eta)}{\eta} \tag{S4}$$

Assuming a constant sequencing error rate $\xi$ per nucleotide, and considering the effect of sequencing error only by substitution, the probability that a sequence will be misidentified as one of its one-mutation neighbors is

$$(1-\xi)^{L-1}\frac{\xi}{3} \tag{S5}$$

A *d*-th order mutant, it has $3L$ neighbors consisting of $d$ (*d*-1)-th mutants (i.e., one of the *d* mutated nucleotides reverted to the wild type), $2d$ *d*-th mutants (i.e., one of the *d* mutated nucleotides changed to one of other two possible mutations), and $3(L-d)$ (*d+1*)-th mutants (i.e., one of the $L-d$ wild type nucleotides mutated). Assuming the real abundance for this *d*-th order mutant is 1, the (*d-1*)-th mutant is $\phi$, and (*d+1*)-th mutant is $1/\phi$. The expected observed abundance for a *d*-th mutant, in a variant pool with mutation rate $\eta$ and sequencing error rate $\xi$ is

$$\rho(d,\,\xi,\,\eta) = (d\phi + 2d + (3L-3d)/\phi)(1-\xi)^{L-1}\frac{\xi}{3} + (1-\xi)^L \tag{S6}$$

The fraction of abundance that originates from its neighbors due to sequencing error is

$$1-\frac{(1-\xi)^L}{\rho(d,\xi,\eta)} \tag{S7}$$

**Figure S15**. Expected error from single-mutation neighbor sequences due to sequencing errors, for different orders of mutants ($d$) and different rates of sequencing error ($\xi$). Family centers ($d = 0$) are the most abundant sequences and would be least affected by sequencing error. With decreased error rate, the fraction of reads resulting from erroneous reads of neighboring sequences is decreased for each order of mutants. The mutation rate in synthesizing the variant pool is 9%.

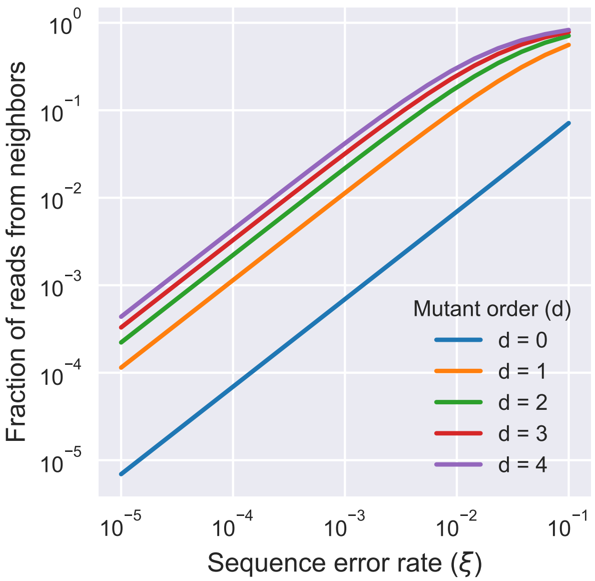

**Figure S16**. Fraction of sequences in which the estimated CI-95 (from bootstrapping or triplicates) includes the true $kA$ values, for sequences with different true $kA$ values. Sequences were ranked by true $kA$ values (from large to small) and each data point indicates the fraction of CI-95 that include the true value in each bin of 25,000 sequences. While CI-95 estimates from bootstrapping consistently includes ~95% of true values, results from triplicates underestimate the uncertainty (i.e., over-confidence), especially for sequences with high $kA$ values.

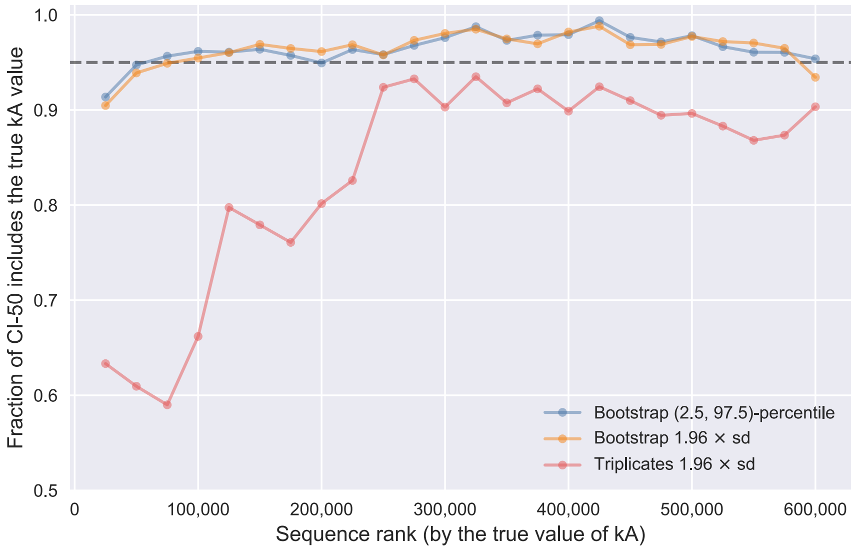

**Figure S17**. Precision (fold-range: 97.5-percentile / 2.5-percentile) did not have a strong dependence on $kA$ (median of bootstrapping samples). A mild decrease to a plateaued was observed as median $kA$ increases. Each dot represents a sequence, colored by Hamming distance $d$ to the family center (see legend).

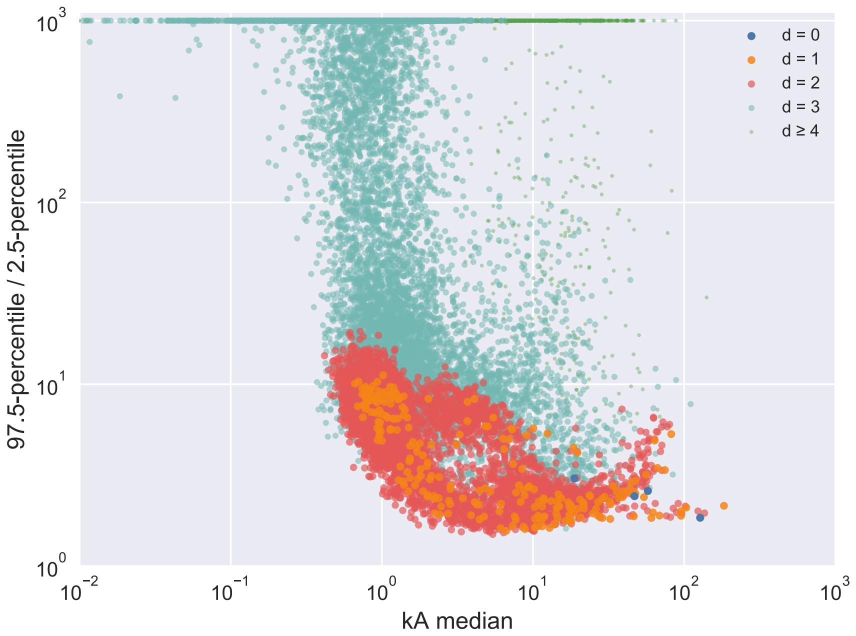

**Figure S18**. Dependence of accuracy (ratio of estimated $kA$ over true $kA$) on counts in the unreacted pool, from simulated count data. The dashed lines correspond to ratios as labeled. Ratios above 100-fold or below 0.01-fold are shown at the borders. The counts of the simulated pool have an uneven distribution similar to the unreacted sample of the variant pool.

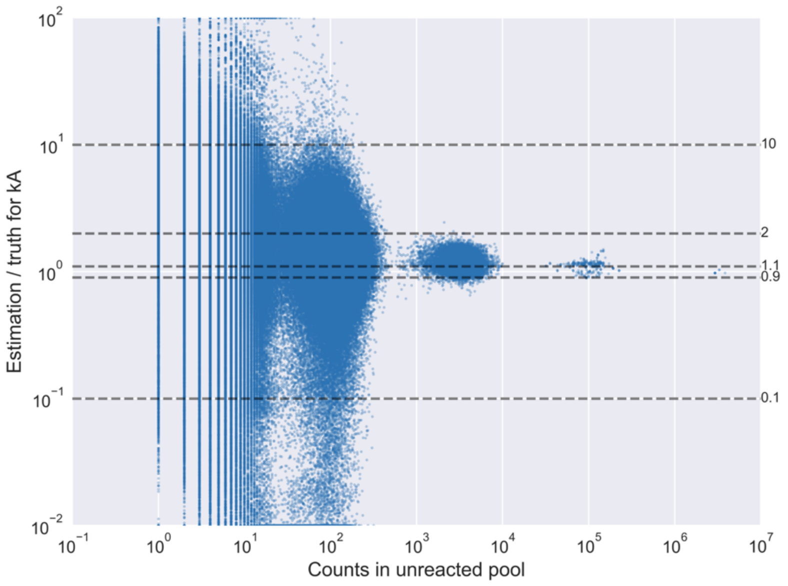

**Figure S19**. Pool evenness for the pool enriched by selection and the designed variant pool. Pool evenness is evaluated by entropy efficiency ($-\sum_{i=1}^{N} p(i) \log p\,(i) \log(N)$, where $p(i)$ is the fraction of sequence $i$ and $N$ is the number of unique sequences) for sequences with various minimum count thresholds in the unreacted samples. As designed, the variant pool is more even (higher entropy efficiency) than the enriched pool.

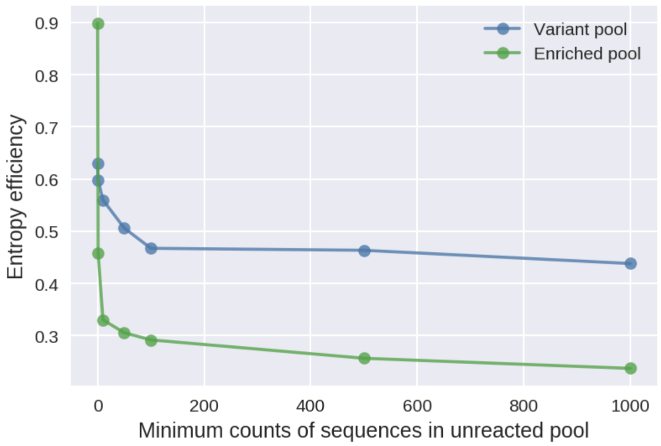

1. Pressman,A.D., Liu,Z., Janzen,E., Blanco,C., Müller,U.F., Joyce,G.F., Pascal,R. and Chen,I.A. (2019) Mapping a Systematic Ribozyme Fitness Landscape Reveals a Frustrated Evolutionary Network for Self-Aminoacylating RNA. *J. Am. Chem. Soc.*, **141**, 6213–6223.
